# Lamina-associated polypeptide 2α is required for intranuclear MRTF-A activity

**DOI:** 10.1101/2021.01.08.425886

**Authors:** Ekaterina Sidorenko, Maria Sokolova, Antti Pennanen, Salla Kyheröinen, Guido Posern, Roland Foisner, Maria K. Vartiainen

**Affiliations:** Institute of Biotechnology, HiLIFE, University of Helsinki, Finland; Institute for Physiological Chemistry, Medical Faculty, Martin Luther University Halle-Wittenberg, Germany; Max Perutz Labs, Center for Medical Biochemistry, Medical University of Vienna, Vienna Biocenter Campus (VBC), Vienna, Austria

## Abstract

Myocardin-related transcription factor A (MRTF-A), a coactivator of serum response factor (SRF), regulates the expression of many cytoskeletal genes in response to cytoplasmic and nuclear actin dynamics. Here we describe a novel mechanism to regulate MRTF-A activity within the nucleus by showing that lamina-associated polypeptide 2α (Lap2α), the nucleoplasmic isoform of Lap2, is a direct binding partner of MRTF-A, and required for the efficient expression of MRTF-A/SRF target genes. Mechanistically, Lap2α is not required for MRTF-A nuclear localization, unlike most other MRTF-A regulators, but is required for binding of MRTF-A to its target genes. This regulatory step takes place prior to MRTF-A chromatin binding, because Lap2α neither interacts with, nor specifically influences active histone marks on MRTF-A/SRF target genes. Phenotypically, Lap2α is required for serum-induced cell migration, and deregulated MRTF-A activity may also contribute to muscle and proliferation phenotypes associated with loss of Lap2α. Our studies therefore add another regulatory layer to the control of MRTF-A-SRF-mediated gene expression, and broaden the role of Lap2α in transcriptional regulation.

## INTRODUCTION

In eukaryotic cells, precise control of gene expression depends on the coordinated work of transcription factors and their cofactors, which also link cellular signaling events to RNA polymerase II recruitment and activation at gene promoters. Many transcription factors themselves have relatively low effector activity, and they therefore cooperate with a cofactor that has this activity (Lambert et al. 2018). One well-described example of such cooperation is serum response factor (SRF), and its cofactors, which together regulate the expression of both immediate-early and many cytoskeletal and muscle-specific genes (Lee et al. 2010; Miano et al. 2007).

Two families of signal-regulated coactivators, which interact with SRF in a competitive manner, mediate differential regulation of SRF target gene expression (Miralles et al. 2003; Zaromytidou et al. 2006). The ternary complex factor (TCF) family of proteins, regulated by the Ras-ERK signaling, mediates the immediate-early transcriptional response (Buchwalter et al. 2004). The myocardin-related transcription factors (MRTFs) MRTF-A (also known as MKL1 or MAL), MRTF-B, and myocardin regulate the expression of muscle-specific and cytoskeletal genes. Expression of myocardin is restricted to smooth and cardiac muscle, but both MRTF-A and MRTF-B are expressed rather ubiquitously, regulate the expression of cytoskeletal genes and respond to the Rho-actin pathway (Miralles et al. 2003; Olson and Nordheim 2010). The subcellular localization and activity of MRTF-A (and MRTF-B) is regulated by cytoplasmic and nuclear actin dynamics, which create a feedback loop, where cytoskeletal dynamics controls the expression of its constituents (Olson and Nordheim 2010). In resting, unstimulated conditions, MRTF-A is mainly localized to the cytoplasm, due to the formation of a complex between monomeric actin and RPEL domain of MRTF-A. The bipartite nuclear localization signal (NLS) of MRTF-A is buried by actin-binding, which restricts import factor access, and thus prevents nuclear import of MRTF-A at high actin monomer concentrations (Mouilleron et al. 2011; Pawlowski et al. 2010; Vartiainen et al. 2007). In addition, actin-binding also promotes nuclear export of MRTF-A via an unknown mechanism (Mouilleron et al. 2011; Vartiainen et al. 2007). Chemical or mechanical stimulation, which induce actin polymerization, for example via activation of the small GTPase RhoA, leads to dissociation of actin monomers from MRTF-A, and consequently increased nuclear import and decreased nuclear export of MRTF-A. This results in nuclear localization of MRTF-A, binding to SRF and expression of MRTF-A/SRF target genes (Pawlowski et al. 2010; Vartiainen et al. 2007).

Also nuclear actin dynamics contribute to MRTF-A regulation. Serum-stimulation leads to formin-dependent polymerization of actin within the nucleus, which is required for MRTF-A nuclear localization and SRF activation (Baarlink et al. 2013). Subsequent study clarified the signaling events leading to nuclear actin polymerization by demonstrating that release of Ca^2+^ from the endoplasmic reticulum targets inner nuclear membrane (INM) localized formin INF2 to promote the formation of linear actin filaments emanating from the INM (Wang et al. 2019). Interestingly, also other proteins localized to the INM have been linked to MRTF-A/SRF regulation. Loss of lamin-A or lamin-A mutations that cause dilated cardiomyopathy-associated laminopathies result in decreased nuclear localization of MRTF-A, and impaired expression of MRTF-A/SRF target genes (Ho et al. 2013). Mechanistically, defects in lamin A cause mislocalization of emerin, an actin-binding protein (Holaska et al. 2004) from the INM to the outer nuclear membrane (ONM), which causes defects in actin dynamics (Ho et al. 2013). In addition, emerin seems to play a critical role in the mechanical regulation of MRTF-A activity especially on stiff substrates (Willer and Carroll 2017). Proteins of the linker of nucleoskeleton and cytoskeleton (LINC) complex, emerin and lamin A are also required for cell spreading-induced nuclear actin polymerization, which depends on integrin activation, and regulates the MRTF-A/SRF pathway (Plessner et al. 2015). LINC complex may also regulate MRTF-A activity by controlling the upstream signaling pathways, since Sun2-containing complexes have been demonstrated to activate RhoA and focal adhesion assembly, while Sun1 antagonizes these activities (Thakar et al. 2017). Taken together, several proteins of the nuclear envelope influence MRTF-A activity, most often by regulating actin dynamics in the cytoplasm or in the nucleus.

This study focuses on the role of lamina-associated polypeptide 2α (Lap2α) as a novel regulator of MRTF-A activity. The Lap2 family contains six alternatively spliced isoforms derived from the *Tmpo* gene (Berger et al. 1996; Harris et al. 1995). All Lap2 isoforms share an N-terminus responsible for chromatin binding, and most of them contain a C-terminal transmembrane domain that anchors them at the INM. These anchored isoforms interact with lamin B, and help to organize heterochromatin at the nuclear periphery (Foisner and Gerace 1993; Lang and Krohne 2003; Dechat et al. 2000). Lap2α is the largest isoform of Lap2 with a unique C-terminus lacking the transmembrane domain and expressed only in mammals (Prufert et al. 2004). Lap2α thus localizes to the nucleoplasm, where it specifically interacts with the nucleoplasmic pool of lamin A affecting lamin A properties and function (Dechat et al. 1998, 2000; Naetar et al. 2008). Lap2α expression, in turn, is affected by Lamin A/C, since the absence of lamin A/C increases Lap2α protein levels (Cohen et al. 2013). Both lamin A/C and Lap2α interact with chromatin, and their binding sites overlap in euchromatic regions. Loss of Lap2α shifts lamin A/C towards more heterochromatic regions, and results in changes in epigenetic histone marks at these sites (Gesson et al. 2016). In addition to general regulation of euchromatin, Lap2α has also been linked to transcription factor regulation. It interacts with the cell cell-cycle regulator protein retinoblastoma (pRb) and affects E2F-pRb-dependent gene expression (Dorner et al. 2006). Indeed, loss of Lap2α leads to inefficient cell-cycle arrest, and hyperproliferation of, for example, erythroid and epidermal progenitor cells in mice (Naetar et al. 2008). Another important role of Lap2α was shown recently in human adipose-derived stem cells (hASCs), where loss of Lap2α suppresses osteogenic differentiation of hASCs via activation of NF-κB transcription factor (Tang et al. 2020). Moreover, loss of Lap2α impairs heart function, and results in deregulation of the cardiac transcription factors GATA4 and MEF2c (Gotic et al. 2010). In addition, both Lap2β and Lap2α regulate the activity of Gli1 transcription factor by regulating its acetylation-dependent trafficking between the nuclear lamina and nucleoplasm (Mirza et al. 2019). Thus, Lap2α appears to affect the activity of different transcriptional factors in various signaling pathways.

In this work, we show that Lap2α is required for efficient MRTF-A/SRF target gene expression, and consequently for cell migration. Unlike other nuclear lamina-associated proteins implicated in this pathway, Lap2α is dispensable for MRTF-A nuclear localization. Mechanistically, Lap2α binds directly to MRTF-A and modulates MRTF-A intranuclear activity prior to chromatin binding.

## RESULTS

### Lap2α is required for MRTF-A-SRF transcriptional activity

Nucleoskeletal proteins were earlier shown to affect MRTF-A nuclear localization, and hence MRTF-A activity, via modulating nuclear actin dynamics (Ho et al. 2013; Kircher et al. 2015; Plessner et al. 2015). We hypothesized that also other nuclear lamina proteins might regulate MRTF-A/SRF pathway, and are here focusing on the role of lamina-associated polypeptide (Lap2) in MRTF-A/SRF regulation. To study this, we first performed siRNA-mediated knock-down of total Lap2, targeting both anchored and nucleoplasmic isoforms (**Fig S1A**), in NIH 3T3 mouse fibroblast cell line, which has been extensively used to study the MRTF-A signaling pathway (Miralles et al. 2003; Rajakyla et al. 2015; Vartiainen et al. 2007). To investigate the effect of Lap2 knockdown on gene expression, we performed RNA sequencing (RNA-seq) analysis of control and Lap2-depleted cells cultivated in serum-starved conditions and after 45 min of serum stimulation **(Fig 1A and Table S1.1**). Serum-responsive genes responded to stimulation in both control and Lap2-depleted cells **(Fig 1A top).** We identified 515 serum-inducible genes (log2FC>1) in control samples and 521 genes that responded to stimulation (log2FC>1) in Lap2-depleted cells (**Fig S1B and Table S1.2**). Of these genes, 356 were common for control and Lap2-depleted samples (**Fig S1B**), but they displayed weaker response to serum stimulation in Lap2-depleted cells compared to control siRNA treated cells **(Fig S1C**). Serum-responsive genes clustered in two groups: the first had increased baseline expression (**Fig 1B top, column 3**) and the second decreased baseline expression and decreased serum-induction (**Fig 1B bottom, columns 3,4**) in Lap2 depleted cells compared to control cells. Out of the 515 genes that were stimulated by serum in control siRNA treated cells, 110 genes displayed decreased baseline expression in starved conditions (n=13), decreased response to serum stimulation (n=68), or both (n=29) (log2FC<0.6) in Lap2-depleted cells **(Table S1.2**). Gene ontology enrichment analysis of these genes revealed a number of actin cytoskeleton–related genes **(Fig 1C and Table S1.3**), strongly indicating that they likely represent MRTF-A/SRF target genes (Morita et al. 2007; Olson and Nordheim 2010).

**Figure 1.**
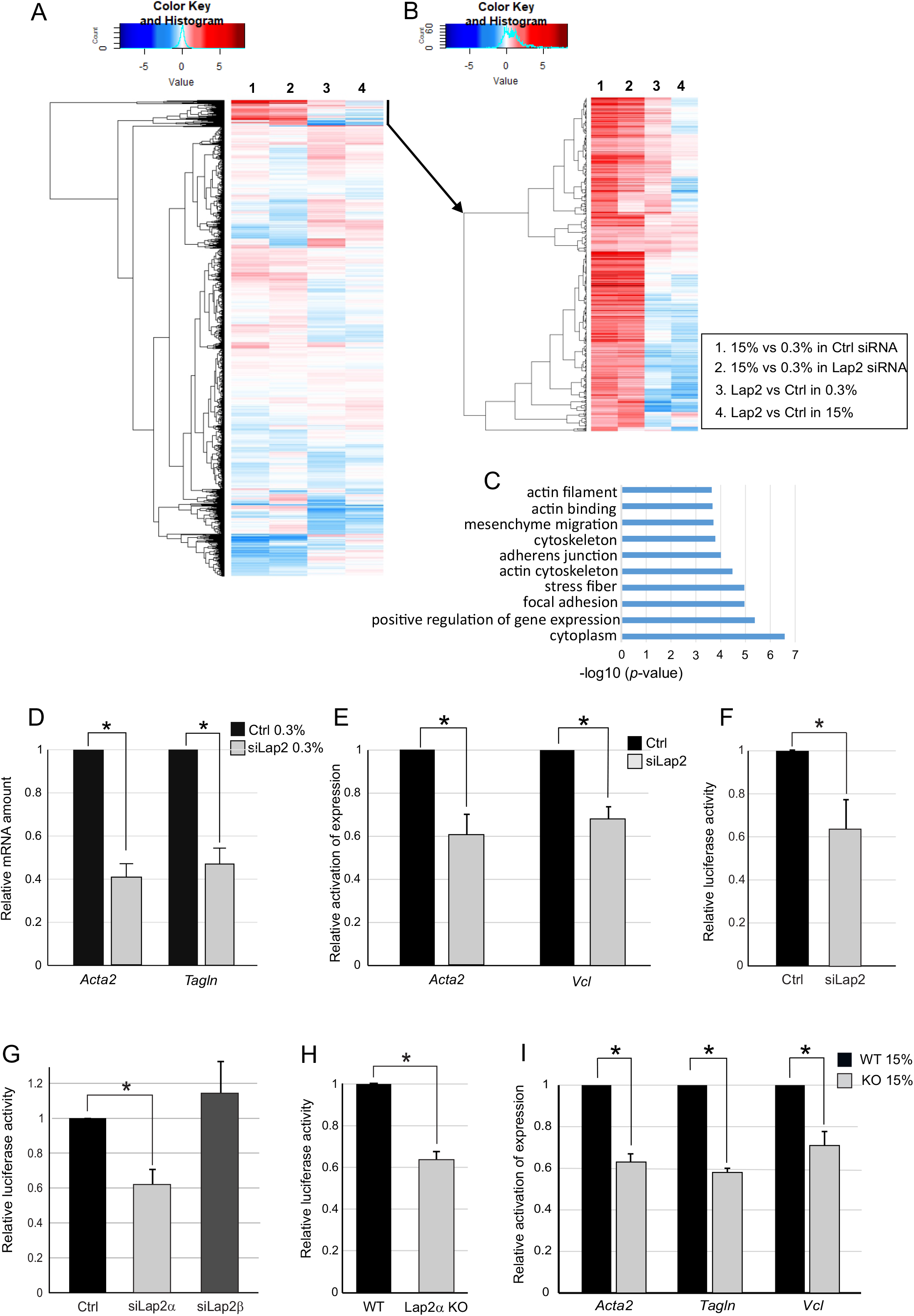
Lap2α is required for MRTF-A/SRF transcriptional activity. **(A)** Heatmap showing expression of all genes in control and Lap2-depleted fibroblasts in serum-starved (0.3% serum) and serum-stimulated (15%) conditions from RNA-seq. **(B)** Heatmap showing expression of serum-responsive genes (subset of panel A) in control and Lap2-depleted fibroblasts in serum-starved and serum-stimulated conditions, as above. **(C)** Gene Ontology enrichment analysis of genes with decreased baseline expression or decreased activation upon serum stimulation in Lap2-depleted NIH 3T3 fibroblasts. **(D)** Depletion of total Lap2 (siLap2) decreases baseline expression (expression in serum starved conditions) of MRTF-A target genes compared to control (Ctrl). Data is the mean with n=5, normalized to Ctrl and error bars are standard error of the mean (s.e.m). Statistically significant differences (*; *P*<0.00001) tested by Student’s *t*-test. *P*-values: Ctrl vs siLap2 for *Acta2* 2.40445E-06, Ctrl vs siLap2 for *Tagln* 9.11726E-05. **(E)** Depletion of total Lap2 decreases transcriptional induction of MRTF-A target genes upon serum stimulation. Data is the mean with n=5, normalized to Ctrl and error bars are s.e.m. Statistically significant differences (*; P<0.001) tested by Student’s *t*-test. *P*-values: Ctrl vs siLap2 for *Acta2* (15%FBS) 0.000513967, Ctrl vs siLap2 for *Vcl* (15% FBS) 0.00044614. **(F)** Depletion of Lap2 decreases serum-induced activation of a SRF reporter gene. Data is the mean with n=5, normalized to Ctrl siRNA and error bars s.e.m. Statistically significant differences (*; *P*<0.01) tested by Student’s *t*-test. *P*-values: Ctrl vs siLap2 (15% FBS) 0.0035 **(G)** Depletion of the -isoform (siLap2α), but not -isoform (siLap2ß) decreases serum-induced SRF reporter activation. Data is mean with n=4, normalized to Ctrl siRNA and error bars are s.e.m. Statistically significant differences (*; *P*<0.01) tested by Student’s *t*-test. *P*-values: Ctrl vs si Lap2α (15% FBS) 0.00455. **(H)** Serum-induced activation of SRF reporter activity is decreased in Lap2α KO mouse dermal fibroblasts (MDF) compared to Lap2a WT MDFs. Data is the mean with n=7, normalized to Lap2α WT MDFs and error bars are s.e.m. Statistically significant differences (*; *P*<0.000001) tested by Student’s t-test. *P*-values: WT vs KO (15% FBS) 6.8425E-07. **(I)** Serum-induced activation of MRTF-A/SRF target gene expression is decreased in Lap2α KO cells (KO) compared to Lap2α WT cells (WT). Data is the mean with n=5, normalized to WT, and error bars are s.e.m. Statistically significant differences (*; *P*<0.001) tested by Student’s t-test. *P*-values: for Acta 3.35E-05, for *Tagln:* 8.21E-07, for *Vcl* 0.000719. See also Supplementary figure 1.

To confirm the recruitment of MRTF-A and SRF to serum-responsive genes, we used chromatin immunoprecipitation (ChIP) followed by deep sequencing (ChIP-seq) in serum-starved and serum-stimulated NIH 3T3 cells. Peak calling (see Materials and Methods for details) revealed that out of the 515 serum-inducible genes identified by RNA-seq, 176 genes contained peaks of either MRTF-A or SRF, or both within 100 kb from the transcription start site (TSS), indicating that most of these genes are direct MRTF-A/SRF targets **(Table S1.2**). Further analysis revealed increased enrichment of both SRF **(Fig S1D)** and MRTF-A **(Fig S1E)** on the SRF containing peaks upon serum stimulation.

To confirm the RNA seq data, we utilized qPCR to study expression of classical MRTF-A/SRF target genes *Acta2*, *Tagln*, and *Vcl.* Depletion of Lap2 resulted in both decreased basal expression (**Fig 1D**), as well as decreased transcriptional induction upon serum stimulation (**Fig 1E**) of these genes. Lap2 depletion also decreased the serum-induced activation of an SRF reporter gene (**Fig 1F**), further suggesting that Lap2 is a potential novel regulator of the MRTF-A/SRF transcriptional activity.

The *Tmpo* gene produces several isoforms of the Lap2 protein (Berger et al. 1996; Harris et al. 1995). To establish which isoform is involved in regulating MRTF-A/SRF activity, we designed siRNAs targeting individually both the nucleoplasmic isoform alpha (α) (ENSMUST00000020123.6) and the anchored isoform β (ENSMUST00000072239.13) (**Fig S1F, G**). Interestingly, depletion of the Lap2α-isoform impaired the activation of the SRF reporter, whereas depletion of Lap2β-isoform had no effect (**Fig 1G**).

To study this further, we took advantage of mouse dermal fibroblasts (MDF) derived from Lap2α knockout mouse (Naetar et al. 2008). As established previously, this cell line does not express any *Lap2α* mRNA (Naetar et al. 2008) or Lap2α protein, whereas expression of Lap2β isoform is unaffected **(Fig S1H)**. Also in this cell model, Lap2α was required for efficient MRTF-A/SRF activity, since both the SRF reporter **(Fig 1H**), and MRTF-A target genes **(Fig 1I**) displayed decreased serum-induced activation in Lap2α knockout (KO) cells compared to Lap2α wild type (WT) cells. These results establish Lap2α as the Lap2 isoform responsible for MRTF-A/SRF regulation.

TCFs are another set of SRF cofactors, which act as general antagonists of MRTF-dependent SRF target gene expression, competing directly with MRTFs for access to SRF (Zaromytidou et al. 2006; Gualdrini et al. 2016). We found that typical TCF/SRF target genes, such as *Egr1, Egr3* and *Fos* were more highly induced by serum in Lap2α KO cells than in Lap2α WT cells (**Fig 2A**). Moreover, we noticed that many immediate-early genes, such as *Ctgf, Ptgs2, Cyr61, Dusp6*, and *Ier3*, were found among genes with high response to serum stimulation and increased baseline expression in Lap2-depleted cells (**Table S1.1**). This result underscores the role of Lap2α in regulating specifically MRTF-A activity, and not SRF in general.

**Figure 2.**
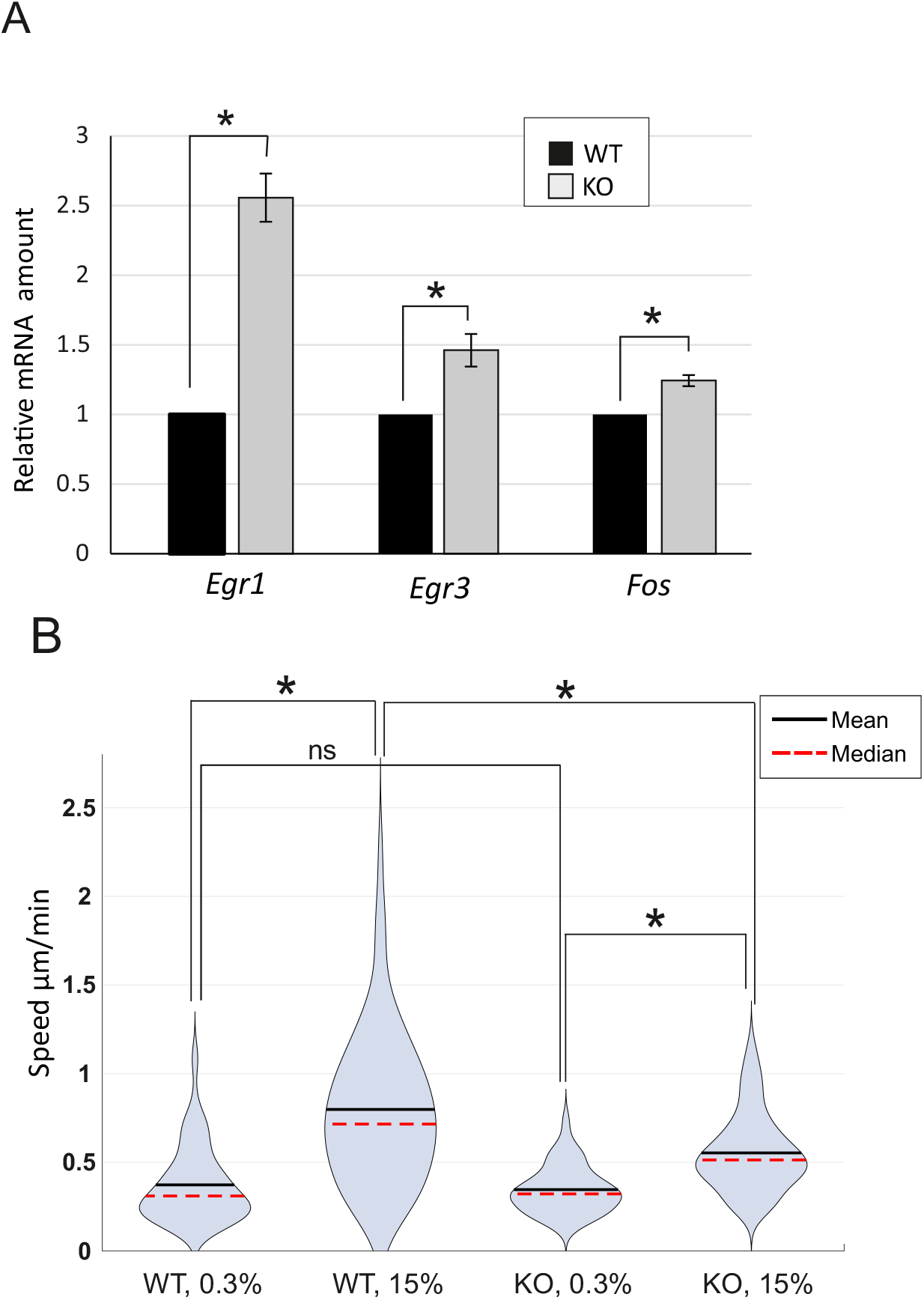
Loss of Lap2α leads to increased expression of TCF/SRF target genes and reduced cell migration. **(A)** Serum-induced activation of TCF/SRF target genes is increased in Lap2α KO MDFs compared to Lap2α WT MDFs. The data is shown as mean with n=4, normalized to Lap2α WT cells and error bars are s.e.m. *Statistically significant differences (*P*<0.01) tested by Student’s *t*-test *P*-values: WT vs KO for *Egr1* 0.000108, WT vs KO for *Egr3* 0.002517, WT vs KO for *Fos* 0.001445 **(B)** Single cell tracking assay shows decreased motility of Lap2α KO MDFs compared to Lap2α WT MDFs in serum-stimulated conditions. Data is presented as violin plots showing distribution of average speeds for each condition with mean and median indicated. Total number of tracks: N = 89 WT 0.3%, N=89 WT 15%; N = 107 KO 0.3%, N=106 KO 15%. Mean values and s.t.d.: WT 0.3% 0.369±0.216; WT 15% 0.783±0.4; KO 0.3% 0.352±0.14; KO 15% 0.556±0.22. Statistically significant differences (*) tested by Kolmogorov-Smirnov test with 0.05 significance level. ns., not significant. D-values: WT 0.3% vs WT 15% 0.614; KO 0.3% vs KO 15% 0.489; WT 15% vs KO 15% 0.383; WT 0.3% vs KO 0.3% 0.174 (non-significant).

To study, if decreased MRTF-A/SRF activation also leads to any phenotypic consequences at the cellular level, we studied cell motility, because many MRTF-A target genes are responsible for cytoskeletal organization, and MRTF-A activity has been shown to be required for efficient cell migration (Leitner et al. 2011; Medjkane et al. 2009; Pipes et al. 2006). Both Lap2α WT and KO cells responded to serum stimulation by increasing their motility (**Fig 2B**). However, in serum-stimulated conditions, the Lap2α KO cells displayed greatly reduced migration speed compared to Lap2α WT cells (**Fig 2B)**. Taken together, we have established that Lap2α is a novel regulator of MRTF-A activity, and required for appropriate expression of MRTF-A target genes, which then play a critical role, for example, in cell migration.

### Lap2α is not required for MRTF-A nuclear localization

One of the main mechanisms to regulate MRTF-A activity takes place via its nucleo-cytoplasmic transport, which depends on actin dynamics both in the cytoplasm and in the nucleus (Baarlink et al. 2013; Vartiainen et al. 2007). Hence, we first investigated whether Lap2α would regulate nuclear localization of MRTF-A, similarly to other components of the nuclear lamina, such as lamin A/C and emerin (Ho et al. 2013). However, endogenous MRTF-A accumulated into the nucleus similarly in Lap2α WT and KO MDFs after 10 min of serum stimulation **(Fig 3A).** Similarly, MRTF-A-GFP did not show any defects in serum-induced nuclear localization upon siRNA-mediated depletion of Lap2α in NIH 3T3 cells **(Fig S3A**). Moreover, MRTF-A nuclear localization upon Leptomycin B treatment, which inactivates Crm1/Exportin1 (Fornerod et al. 1997), the established nuclear export receptor for MRTF-A (Pawlowski et al. 2010; Vartiainen et al. 2007) or upon Cytochalasin D treatment, which directly disrupts the MRTF-A-actin complex (Vartiainen et al. 2007) was not perturbed in Lap2α KO cells (**Fig S3B,C**). These results indicate that nuclear localization of MRTF-A is not dependent on Lap2α. To our surprise, we found that in Lap2α KO fibroblasts, MRTF-A was present in nuclei of many cells even in starved conditions **(Fig 3A, B, C).** This indicates that unlike lamin A/C and emerin (Ho et al. 2013), Lap2α is not required for MRTF-A nuclear localization, but rather modulates its activity within the nucleus.

**Figure 3.**
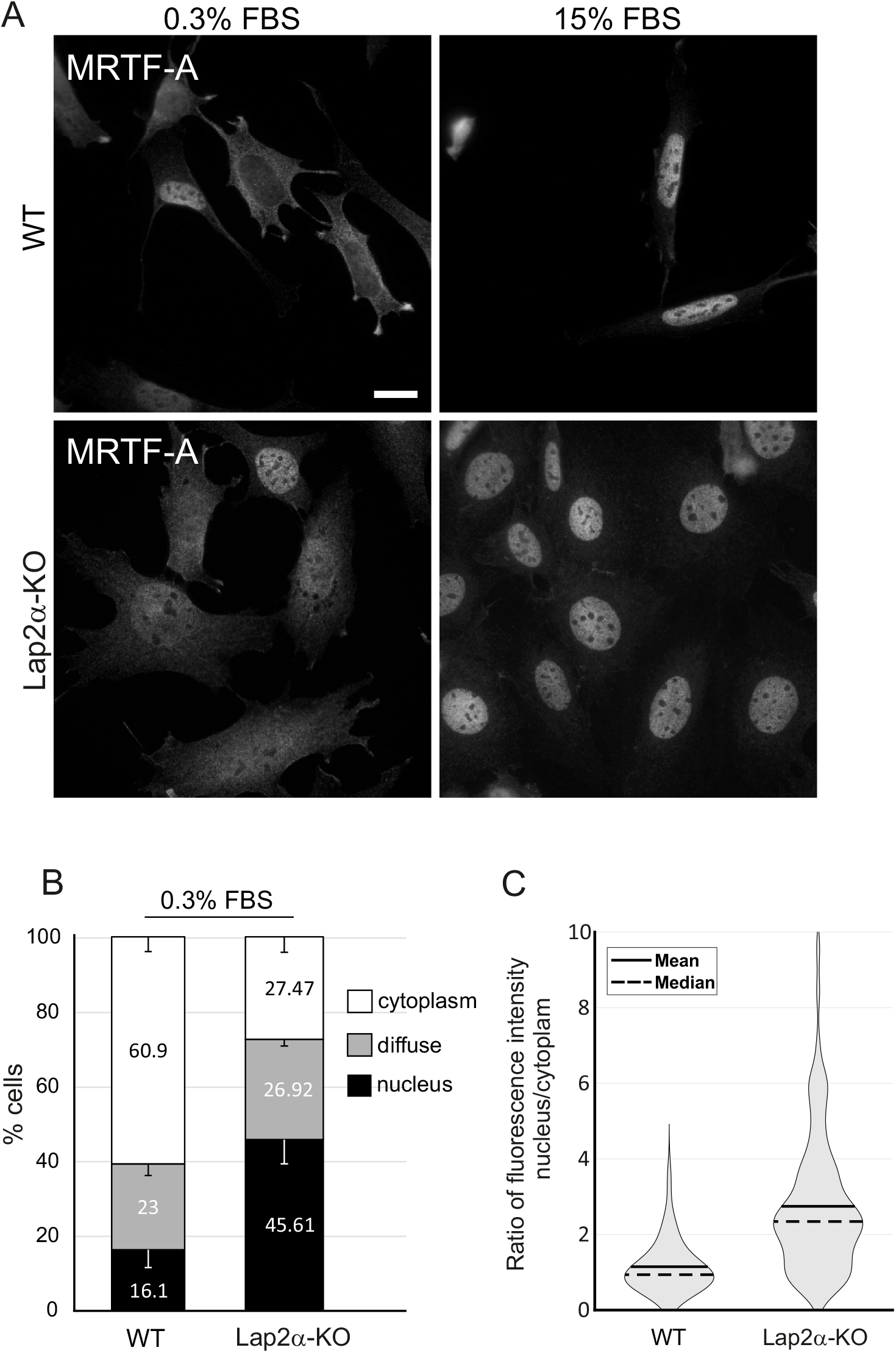
Lap2α is not required for MRTF-A nuclear localization. **(A)** MRTF-A localization in Lap2α WT and KO MDFs in serum starved (0.3% FBS) cells and after stimulation 10 min stimulation with 15% FBS. scale bar = 20 um **(B)** Quantification of MRTF-A localization in Lap2α WT and KO MDFs in serum-starved conditions (0.3%). n=2; error bars are s.d. 100-190 cells per point. **(C)** Fluorescence intensity ratio between the nucleus and the cytoplasm for endogenous MRTF-A in WT and Lap2a KO MDF cells under starved conditions (0.3% FBS). Violin plot showing distribution of cells according to the intensity ratio (nuc/cyt). For WT: n=117, mean=1.1471, median= 0.9364, s.d.= 0.639573. For KO: n=92, mean=2.7451, median=2.3436, s.d.=1.763705. See also S3.

### Lap2a is required for MRTF-A recruitment to SRF target genes

To further study the role of Lap2α in the regulation of MRTF-A activity, we examined MRTF-A-chromatin interaction in Lap2α WT and KO MDFs. We performed chromatin immunoprecipitation (ChIP) followed by deep sequencing (ChIP-seq) of MRTF-A in MDFs under serum starved (0.3% FBS) condition and 45 min after serum stimulation (15% FBS). In addition, Pol II phosphorylated at serine 5 (Pol II S5P) was included to monitor transcription activation of serum-inducible genes.

First, based on peak calling we identified the total number of MRTF-A peaks in Lap2α WT and KO cells under both conditions (**Fig 4A and S4A, Table S2**). Analysis with MASC2 identified 1369 MRTF-A peaks in WT cells and 611 peaks in KO cells in serum stimulated conditions. Of these peaks, 442 were common for Lap2α WT and KO cells **(Fig 4A).** As expected, in WT MDF cells, the number of MRTF-A peaks increased after serum stimulation more than two times (556 in 0.3% vs 1369 in 15% FBS). Surprisingly, in Lap2α KO MDFs, serum stimulation reduced the number of MRTF-A peaks (1004 in 0.3% vs 611 in 15% FBS). Total number of MRTF-A peaks in starved conditions was also higher in KO cells compared to WT cells (**Fig S4A, Table S2**). This may be due to increased nuclear localization of MRTF-A in KO cells compared to WT cells (**Fig 3).**

**Figure 4.**
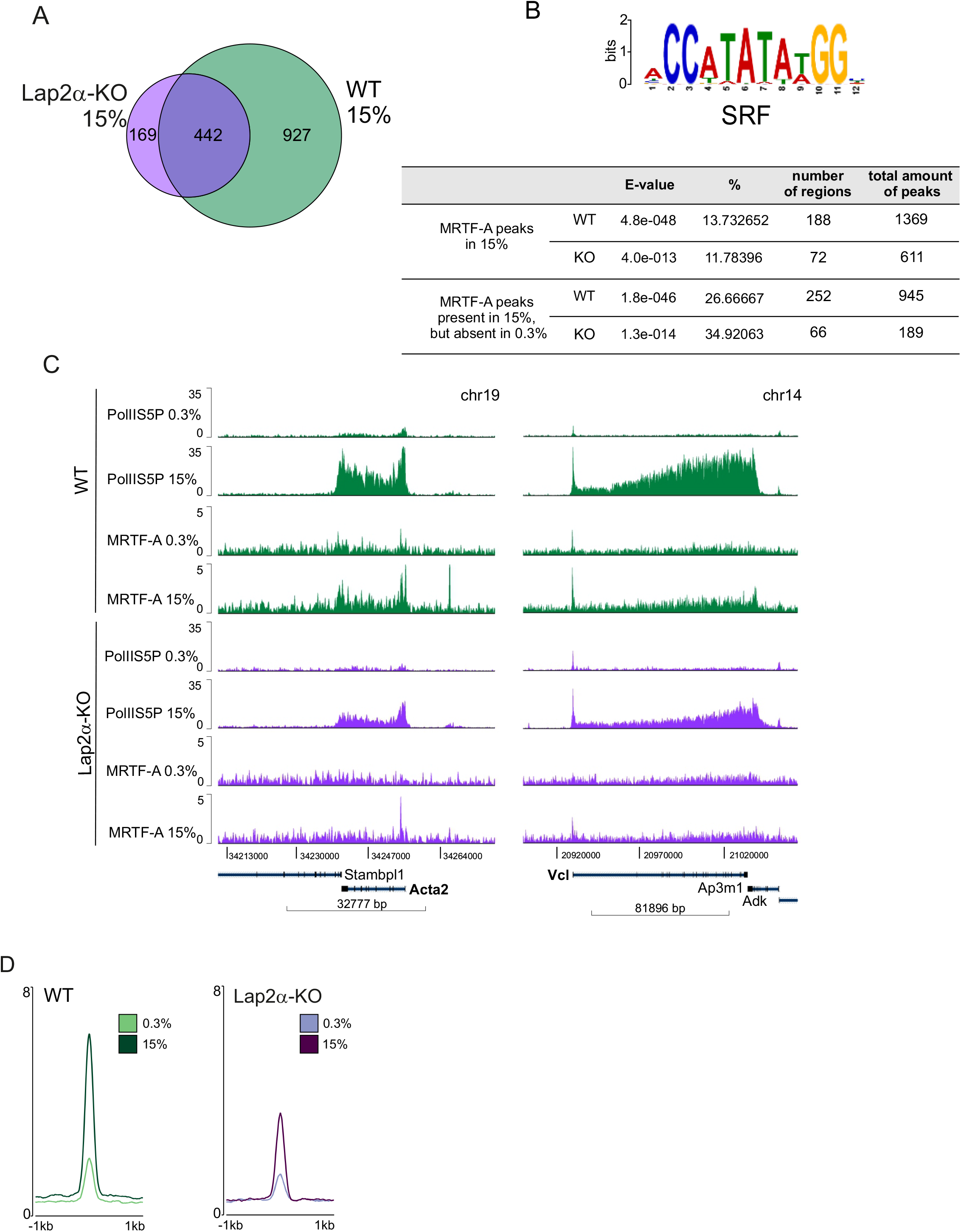
Lap2a is required for MRTF-A recruitment to SRF target genes. **(A)** Venn diagram showing overlap of MRTF-A peaks in WT and Lap2α KO cells under serum-stimulated (15%) conditions from chromatin immunoprecipitation followed by deep-sequencing (ChIP-seq). **(B)** MEME-ChIP analysis identifies SRF motif (upper part) within the MRTF-A peaks found in 15% FBS (upper table) and peaks present in 15% FBS but absent in 0.3% FBS (lower table). **(C)** Binding of Pol II S5P and MRTF-A on MRTF-A target genes *Acta2* and *Vcl* in Lap2α WT and KO MDFs in serum-starved (0,3%) and stimulated (15%) conditions. **(D)** Coverage of MRTF-A on the peaks with SRF motif in Lap2α WT vs KO cells in serum-starved (0,3%) and serum-stimulated (15%) conditions.

We used MEME-ChIP for motif enrichment analysis and found that, as expected, the MRTF-A peaks detected in serum-stimulated conditions were significantly enriched for the SRF motif in both Lap2α WT and KO cells **(Fig 4B**). Interestingly, the coverage of MRTF-A on these SRF motif-containing peaks (**Table S3.1**) was much less in Lap2α KO cells than in Lap2α WT cells in serum stimulated conditions **(Fig 4C,D).** This indicates that Lap2α is required for MRTF-A recruitment to SRF target genes. According to the MEME-ChIP data analysis, MRTF-A peaks were, in addition to SRF, also enriched for motifs of several other transcription factors (TF) in both WT and Lap2α KO cells (**Table S4).** This data indicates that MRTF-A can participate also in SRF-independent transcriptional regulation, as suggested earlier (Gurbuz et al, 2014).

For further analysis, we chose 36 genes, which contained MRTF-A peaks with an SRF binding motif on their promoters (less than 1 kb from TSS) (**Table S3.2)**. This group of genes included many well-established targets of the RhoA/MRTF-A/SRF pathway such as *Acta2*, *Vcl* **(Fig 4C)**, *Myhl9, Cap1, Myo1e*, and *Tuba1c*. On these genes, both MRTF-A binding to the SRF motif, and the amount of Pol II S5P on the gene body, were decreased in Lap2α KO cells compared to Lap2α WT cells **(Fig 4D**), corroborating the decreased expression of these genes in Lap2α KO cells as measured by RT-qPCR (**Fig 1I**).

Thus far, our data indicates that Lap2α is required for efficient binding of MRTF-A to MRTF-A/SRF target genes, and consequently needed for their expression. Since Lap2α can also interact with chromatin, and has been shown to bind especially euchromatic regions (Gesson et al. 2016), we next tested whether Lap2α would also interact with MRTF-A/SRF target genes. However, our ChIP-seq analysis failed to detect any enrichment of Lap2α on MRTF-A peaks either in Lap2a WT or in KO cells (**Fig 5A,B)**, strongly suggesting that Lap2α does not regulate MRTF-A activity at the promoter level. Indeed, previous studies have indicated that Lap2α, in complex with nucleoplasmic Lamin A/C, may be involved in the establishment of euchromatin environment via epigenetic mechanisms rather than regulating gene expression by directly binding to promoters (Gesson et al. 2016). To study if Lap2α modulates the epigenetic landscape of also MRTF-A/SRF target genes, we analyzed the levels of the active H3K4me3 and H3K9Ac histone marks on the promoter region of the 36 MRTF-A peaks (**Table S3.2,** see also **Fig S4B,C** for Pol II S5 and MRTF-A enrichment on these genes), and on all expressed genes in Lap2α WT and KO cells (**Fig 5C, D**; ChIP-seq data from (Gesson et al. 2016)). Although the MRTF-A/SRF target genes displayed decreased amount of RNA Pol II S5P in Lap2α KO cells compared to WT cells (**Fig S4B**), the levels of H3K4me3 and H3K9Ac were very similar in both cell lines **(Fig 5C**). In contrast, genes that were previously shown to display altered expression in Lap2α KO cells displayed increased and decreased levels of H3K4me3 and H3K9Ac histone marks on up-regulated and down-regulated genes, respectively (**Fig S5A-D**: data from Gesson et al, 2016). Although H3K9Ac levels on the selected 36 genes appeared slightly higher in Lap2α WT than in Lap2α KO cells (**Fig 5C, right panel**), the same difference remained for all expressed genes (**Fig 5D right panel**), indicating that it was not a specific characteristic of MRTF-A/SRF target genes. Taken together, Lap2α does not show any preferential binding to MRTF-A/SRF target genes, and therefore the influence on H3K4me3 and H3K9Ac on these genes does not extend beyond the overall effect that Lap2α has on these histone marks.

**Figure 5.**
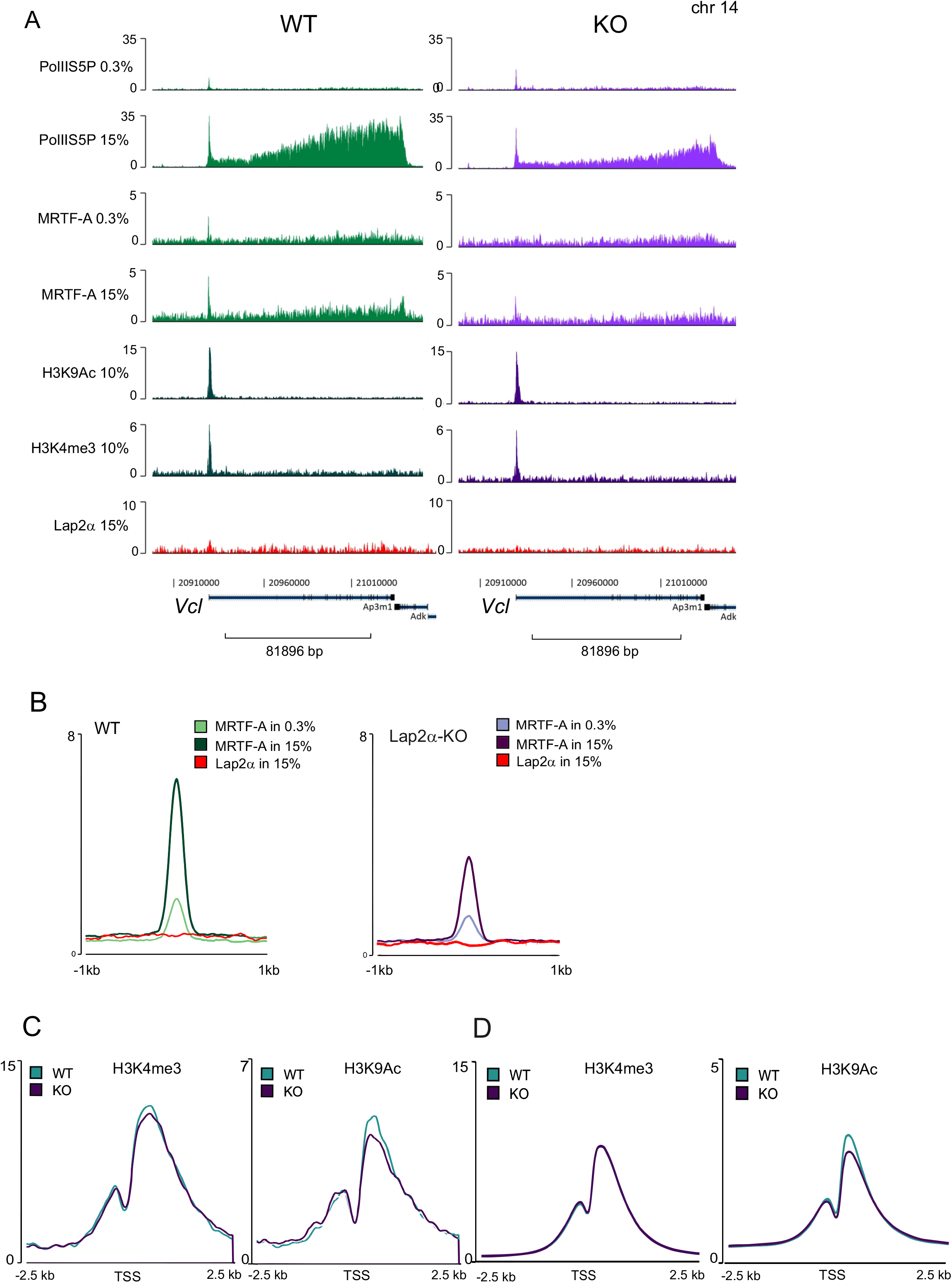
Lap2α neither interacts with MRTF-A/SRF target genes nor modulates their active histone marks. **(A)** Binding of Pol II S5P, MRTF-A, H3K9Ac, H3K4me3 and Lap2α by ChIP-seq to MRTF-A target gene *Vcl* in Lap2α WT and KO fibroblasts under serum starved (0,3%) and serum stimulated (15%) conditions. Data for H3K9Ac and H3K4me3 binding were used from Gesson et al, 2016 **(B)** Coverage of Lap2α and MRTF-A on MRTF-A peaks with SRF motif in Lap2α WT and KO MDFs cells in serum-starved (0.3%) and serum-stimulated (15%) conditions. **(C)** Histone modifications at the TSS of 36 selected genes (genes that contained MRTF-A peaks with an SRF-binding motif on their promoters, Table **S3.2**) in WT and KO MDF. Left: H3K4me3; right: H3K9Ac **(D)** Histone modifications at the TSS of all expressed genes in WT and KO MDF. Left: H3K4me3; right: H3K9Ac

### MRTF-A binds to Lap2α

Our data so far has demonstrated that Lap2α is required for the proper recruitment of MRTF-A to its target genes. To explore the mechanism of this regulation, we hypothesized that Lap2α might interact with MRTF-A, and thus affect its activity. To study this, we performed co-immunoprecipitation (co-IP) studies in NIH 3T3 fibroblasts. First, the interaction between full-length proteins was confirmed. Antibody against tagged MRTF-A efficiently precipitated endogenous and tagged Lap2α (**Fig 6 A, B**) and tagged Lap2α precipitated tagged full-length MRTF-A **(Fig 6C)**. However, we did not detect any interaction between Lap2α and SRF in similar co-immunoprecipitation experiments (**Fig S6)**, further confirming our results that Lap2α is specifically involved in regulating MRTF-A, and not SRF in general (**Fig 2A**).

**Figure 6.**
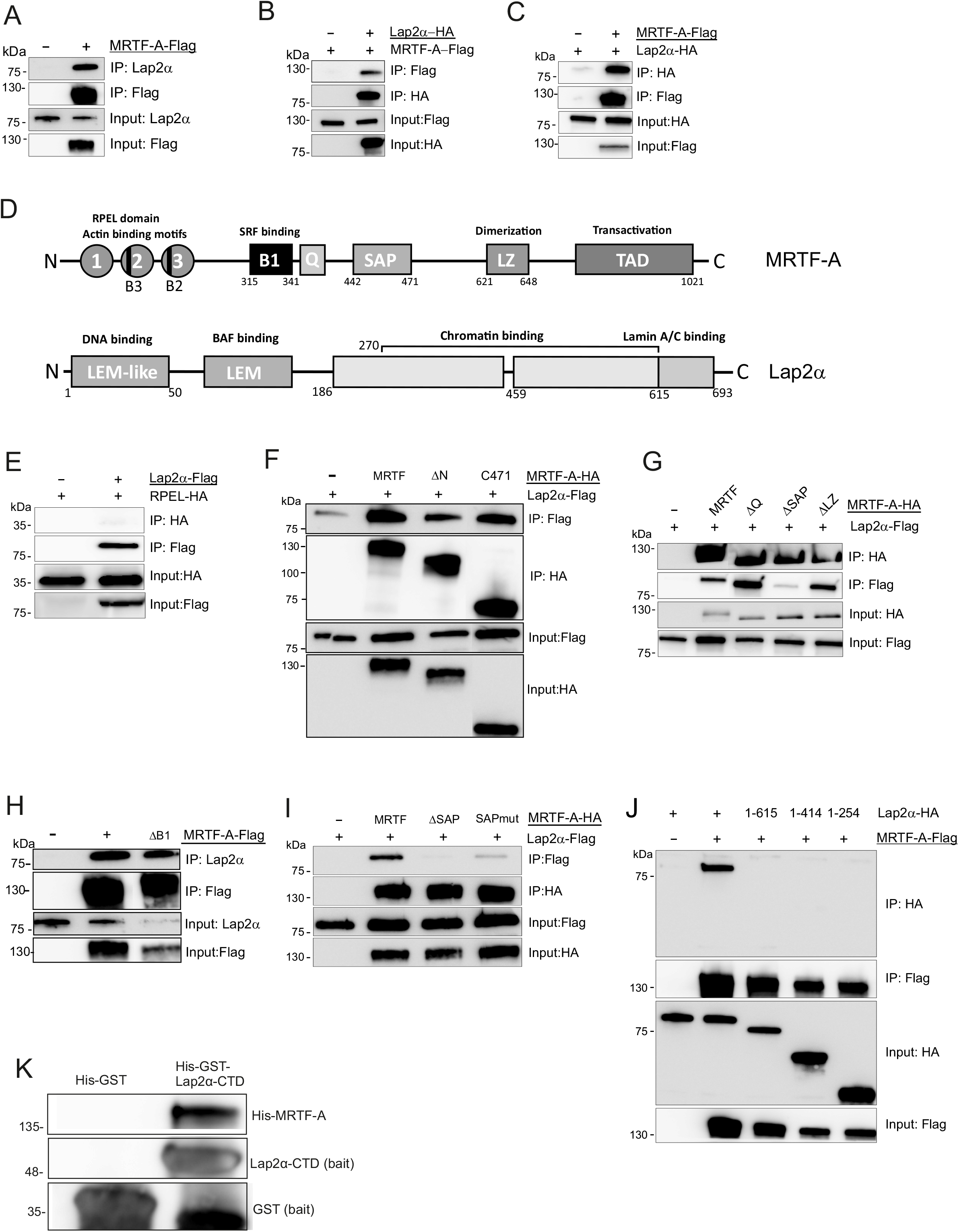
MRTF-A binds to Lap2α. **(A)** Flag-tagged MRTF-A co-immunoprecipitates endogenous Lap2α. Transfected constructs are shown above the blots with the construct used as a bait underlined, protein weight markers on the left and the sample (immunoprecipitate or input) with the antibody used in Western on the right. **(B)** HA-tagged Lap2α co-immunoprecipitates MRTF-A-Flag; data shown as in A. **(C)** Flag-tagged MRTF-Aco-immunoprecipitates Lap2α-HA; data shown as in A. **(D)** Schematic presentation of MRTF-Aand Lap2α domain structures. MRTF-Adomains: 1,2,3 - three RPEL motifs with two basic domains (B3 and B2); B1-basic domain responsible for SRF binding; Q-glutamine-rich domain; SAP-SAP (SAF A/B-Acinus-PIAS)-domain; LZ - leucine-zipper domain; TAD-transcription activation domain. Lap2α domains: LEM-like, LEM (Lap2-Emerin-MAN1 domain); chromatin binding domain; Lamin A/C binding domain. **(E)** Actin-binding RPEL domain of MRTF-A is not involved in the interaction with Lap2α; data shown as in A. **(F)** N- and C-terminus of MRTF-Aare not involved in the interaction with Lap2α; data shown as in A. **(G)** Deletion of the SAP domain (ΔSAP), but not of the Q (ΔQ) and LZ (ΔLZ) domains MRTF-A reduces the interaction with Lap2α, data shown as in A. **(H)** SRF-binding B1-domain of MRTF-A is not involved in the interaction with Lap2α, data shown as in A. **(I)** Introducing point mutations into the SAP domain of MRTF-A impairs binding with Lap2α, data shown as in A. **(J)** Deleting the C-terminus of Lap2α impairs co-immunoprecipitation with MRTF-A, data shown as in A. **(K)** Pulldown assay of His-MRTF-A using His-GST-Lap2α-CTD (C-terminal domain) or His-GST as baits. All proteins were detected with anti-His antibody.

Next, we utilized different deletion constructs of MRTF-A to identify the functional domain (**Fig 6D**) responsible for the Lap2α interaction. First, we tested a construct consisting of the N-terminal RPEL domain of MRTF-A, which is responsible for actin-binding and involved in MRTF-A nuclear import (Guettler et al. 2008; Pawlowski et al. 2010). However, no interaction was detected (**Fig 6E)**, which goes in line with the experiments showing that Lap2α does not regulate MRTF-A nuclear import (**Fig 3**). Also MRTF-A-ΔN, which lacks the N-terminus including the RPEL domain, efficiently co-precipitated Lap2α, further confirming that Lap2α binding takes place beyond the RPEL domain (**Fig 6F**). Another MRTF-A deletion construct without C-terminus (MRTF-A C471) also bound to Lap2α (**Fig 6F)**. Further experiments with other MRTF-A deletion constructs indicated that while the B1 domain (responsible for the SRF binding), leucine zipper (LZ) and Q domains were not required for efficient Lap2α interaction, the construct with deletion of the SAP domain (ΔSAP) showed greatly reduced interaction with Lap2α **(Fig 6G, H**). Because deletion of the whole domain might disrupt protein structure and thus affect its binding ability, we introduced point mutations inside the SAP domain (see Materials and methods for details). Similarly to the deletion of the SAP domain, also these point mutations impaired the interaction between MRTF-A and Lap2α **(Fig 6I**), confirming the role of the MRTF-A SAP domain in Lap2α binding. Next we utilized C-terminal truncations of Lap2α (**Fig 6D**) to delineate the MRTF-A binding site. Unfortunately, N-terminal truncations of Lap2α were not expressed sufficiently for co-IP experiments, as has been reported also earlier (Vlcek et al. 1999). Nevertheless, Lap2α constructs with deletions of the C-terminus (1-615, 1-414, 1-254) displayed similar expression levels to the full-length protein, but did not show any interaction with MRTF-A **(Fig 6J).** These results demonstrate that MRTF-A and Lap2α interact in cells, and that this interaction takes place via SAP-domain of MRTF-A, and requires an intact C-terminus of Lap2α.

To study whether MRTF-A binds directly to Lap2α, we performed a pull-down assay with MRTF-A expressed and purified from insect cells, and recombinant His-GST-tagged C-terminal domain (CTD) of Lap2α expressed in bacterial cells. MRTF-A was detected on beads coated with Lap2α-CTD, but not on beads with GST **(Fig 6K),** demonstrating a direct interaction between MRTF-A and C-terminus of Lap2α. This result also supported our Co-IP data about the role of Lap2α C-terminus in the interaction with MRTF-A.

## Discussion

The nucleoskeleton – a “dynamic network of networks”-provides not only mechanical support to the nucleus but contributes significantly to many nuclear activities, such as expression of specific sets of genes (Mattout-Drubezki and Gruenbaum 2003; Simon and Wilson 2011). Previous studies have demonstrated that many components of the nucleoskeleton, especially nuclear actin and several components of the nuclear lamina, such lamin A/C, emerin and the LINC complex (Baarlink et al. 2013; Ho et al. 2013; Willer and Carroll 2017), regulate the activity of MRTF-A, a transcription coactivator of SRF. Here we extend these studies by showing that Lap2α, the nucleoplasmic isoform of Lap2, is a novel regulator, and a direct binding partner, of MRTF-A, and required for the expression of MRTF-A-SRF target genes by modulating MRTF-A chromatin binding. Our studies therefore add another regulatory layer to the control of MRTF-A-SRF-mediated gene expression, and broaden the role of Lap2α in transcriptional regulation.

The *Tmpo* gene produces Lap2 proteins with very different functional properties. The anchored isoforms, including the Lap2β isoform, contain transmembrane domains, and are therefore important components of the nuclear envelope, interact with lamin B and help to organize heterochromatin at the nuclear periphery (Berger et al. 1996; Dechat et al. 2000). In terms of gene expression, these anchored isoforms are mostly involved in gene silencing. For example, Lap2β has been shown to influence repressive epigenetic modifications via interaction with HDAC3, and to reduce transcriptional activity of the E2F-DP complex (Nili et al. 2001; Somech et al. 2005). In contrast, Lap2α lacks the transmembrane domain and localizes to the nucleoplasm, where it associates with euchromatin (Gesson et al. 2016). Our experiments with siRNAs targeting specifically the anchored and nucleoplasmic isoforms of Lap2 (**Fig S1D, E**), clearly demonstrate that Lap2α is required for MRTF-A-SRF activity, whereas the anchored isoforms are dispensable (**Fig 1G**). Further studies utilizing the MDF cell line derived from Lap2α KO mice (Naetar et al. 2008) confirmed the importance of Lap2α for MRTF-A/SRF function (**Fig 1H, I**). The specificity of Lap2α isoform in regulating MRTF-A is also supported by our binding experiments, which showed that an intact C-terminus, which is unique to Lap2α, is required for binding to MRTF-A (**Fig 6J, K**).

One of the main mechanisms to control MRTF-A activity is via its nucleo-cytoplasmic shuttling, which is regulated by both cytoplasmic and nuclear actin dynamics (Baarlink et al. 2013; Vartiainen et al. 2007). Indeed, several nucleoskeletal proteins including lamin A/C and emerin have been shown to regulate MRTF-A activity by affecting its nuclear accumulation. These proteins are required for the proper polymerization of nuclear actin, and consequently for MRTF-A activation (Ho et al. 2013; Plessner et al. 2015). However, we did not observe any decrease in MRTF-A nuclear localization in Lap2α KO cells, or upon siRNA-mediated silencing of Lap2α, in any conditions that induce MRTF-A nuclear accumulation (**Fig 3A; S3 A,B,C**). Interestingly, in Lap2α KO cells, many cells contained MRTF-A in the nucleus already in starved conditions **(Fig 3B**). This could reflect activation of a feedback loop, where defective expression of many cytoskeletal genes leads to imbalance in actin polymerization, and consequently to nuclear localization of MRTF-A to correct the transcriptional defects.

The mechanism by which Lap2α impinges on MRTF-A activity is therefore different than the other nucleoskeletal proteins utilize. The fact that in Lap2α KO cells MRTF-A can localize to the nucleus (**Fig 3A**), but target gene expression is impaired (**Fig 1I**), implies that the regulation impinges on nuclear MRTF-A. Indeed, ChIP-seq studies revealed reduced binding of MRTF-A to SRF target genes in Lap2α KO cells (**Fig 4C, D**). Also Lap2α is a chromatin-binding protein, containing LEM and LEM-like domains at the N-terminus, which mediate chromatin binding via Barrier-to-Autointegration Factor (BANF1) and by directly binding to DNA, respectively (Cai et al. 2001; Laguri et al. 2001; Segura-Totten and Wilson 2004; Vlcek et al. 1999). In addition, the C-terminal region of Lap2α associates even with mitotic chromosomes (Vlcek et al. 2002). Furthermore, genome-wide binding studies have indicated that Lap2α associates, together with A-type lamins, with large euchromatic regions (Gesson et al. 2016). However, we failed to detect any clear overlap between MRTF-A and Lap2α enrichment on MRTF-A/SRF target genes (**Fig 5B**), indicating that Lap2α and MRTF-A are not interacting with target gene promoters as a complex.

The exact mechanism by which Lap2α regulates MRTF-A activity awaits further studies. One putative mechanism could be via regulation of the epigenetic landscape of MRTF-A/SRF target genes. Indeed, previous studies had reported altered levels of active histone marks H3K9Ac and H3K4me3 on genes that displayed differential expression in Lap2α KO cells (Gesson et al. 2016; **Fig S5A-D**). However, at least these histone marks were not specifically altered on MRTF-A/SRF target genes, when compared to all transcribed genes (**Fig 5C**), further highlighting the inability of Lap2α to enrich at MRTF-A/SRF target genes. However, we cannot exclude the possibility that these minor changes in H3K9Ac and H3K4me3, or in other Lap2α-dependent histone modifications, or other Lap2α-mediated effects on chromatin environment could influence the ability of MRTF-A to bind to its target genes. Mainly *in vitro* studies have shown that MRTF-A/SRF interaction is favored at sites, where the DNA is easily bent (Zaromytidou et al. 2006). Perhaps this property makes MRTF-A exquisitely sensitive to the chromatin environment, whereas the TCF/SRF interaction, occurring preferentially on a defined Ets-motif (Treisman et al. 1992), would be less sensitive. Nevertheless, since our co-immunoprecipitation and pull-down experiments demonstrate a physical interaction between Lap2α and MRTF-A, but not SRF (**Fig 6, S6**), we favor a model, where Lap2α regulates MRTF-A activity prior to chromatin-binding. This could take place via post-translational regulation of MRTF-A. For example, MRTF-A is phosphorylated on multiple residues, and phosphorylation both positively and negatively regulates MRTF-A activity (Panayiotou et al. 2016). On the other hand, UBC9-mediated SUMOylation of MRTF-A reduces its transcriptional activity (Nakagawa and Kuzumaki 2005). Another potentially interesting post-translational modification in this context is acetylation, since recent studies have shown that it can both positively and negatively influence MRTF-A activity. SIRT1-mediated deacetylation enhances the transcriptional activity of MRTF-A, and promotes its recruitment to the collagen type I promoters (Yang et al. 2020). However, in monocytes, acetylation by PCAF promotes nuclear translocation of MRTF-A and consequent interaction with transcription factor NF-κB, thus affecting pro-inflammatory transcription (Yu et al. 2017). MRTF-A also interacts with HDAC6, and its inhibition causes increased acetylation and levels of MRTF-A, leading to enhanced transcriptional activity of SRF. Since HDAC6 is a cytoplasmic protein, this regulation likely takes place prior to MRTF-A nuclear accumulation (Zhang, M. et al. 2018). On the other hand, Lap2 isoforms have been shown, through competitive interactions, to regulate the activity of Hedgehog transcription factor Gli1 by controlling its acetylation, and thereby its movement between the nuclear lamina and nucleoplasm. The nuclear membrane anchored Lap2β maintains a reserve of Gli1 at the INM in acetylation-dependent manner, while Lap2α, promoted by aPKC, scaffolds Gli1 to HDAC1 for deacetylation and release from the INM (Mirza et al. 2019). In the future, it will be interesting to study, whether the direct interaction between MRTF-A and Lap2α facilitates MRTF-A activity via control of its acetylation.

Decreased MRTF-A/SRF activity upon lack of Lap2α revealed interesting aspects and possibilities into both MRTF-A function and phenotypical effects of Lap2α. In line with the literature describing the competitive interactions between MRTFs and TCFs for regulating SRF-mediated transcription (Zaromytidou et al. 2006), we observed increased activation of several canonical TCF target genes, such as *Egr1, Egr3* and *Fos* in Lap2α KO cells compared to Lap2α WT cells (**Fig 2A**). We thus propose that the absence of Lap2α shifts the balance between MRTF and TCF-dependent gene expression towards TCFs, promoting a proliferative program. This could therefore contribute, together with the regulation of pRb, to the hyperproliferative phenotype observed upon loss of Lap2α in many contexts (Dorner et al. 2006; Naetar et al. 2008). Analysis of Lap2α-deficient mice have also revealed phenotypes associated with both skeletal and heart muscle function. Loss of Lap2α leads to an increase in myofiber-associated stem cell pool and shifts the myofiber-type ratio towards fast fiber types (Gotic et al. 2010). Moreover, loss of Lap2α leads to systolic dysfunction in younger mice, as well as increased susceptibility for fibrosis in old mice. These defects in heart function are accompanied by deregulation of GATA4 and Mef2c transcription factors (Gotic et al. 2010). It is tempting to speculate that the novel function described here for Lap2α in MRTF-A regulation might play a role in muscle tissues as well. Indeed, many MRTF-A/SRF target genes are muscle-specific genes (Mack and Hinson 2005; Olson and Nordheim 2010; Selvaraj and Prywes 2003) and MRTFs in general are considered as important transcription cofactors for both the development of the cardiovascular system and its adaptation to injury and stress (Parmacek 2007). MRTF-A is also required for skeletal muscle differentiation both *in vivo* and *in vitro* (Cenik et al. 2016; Selvaraj and Prywes 2003). MRTF-A is tightly regulated in skeletal muscle, and increased expression levels have been reported upon regeneration of skeletal muscle injuries (Mokalled and Poss 2018), although decrease in MRTF-A levels are needed during myoblast fusion (Charrasse et al. 2006; Iwasaki et al. 2008). One mechanism to control MRTF-A abundance during myogenic differentiation is by microRNAs miR24-3p and miR-486-5p, which bind to the 3’UTR of the MRTF-A transcript and reduce MRTF-A expression (Holstein et al. 2020). Deregulation of MRTF-A/SRF pathway has been described in mice and cells expressing dilated cardiomyopathy (DCM)-causing lamin A/C mutations (Ho et al. 2013). Interestingly, a mutation in Lap2α that lowers its affinity for lamin A/C has been identified in a family suffering from DCM (Taylor et al. 2005). It would be interesting to study whether MRTF-A/SRF target gene expression would be affected also in this DCM context. Further studies are needed to elucidate the functional significance of MRTF-A-mediated transcription to the multiple phenotypes associated with loss of Lap2α, and its potential significance to diseases caused by mutations in either Lap2α or lamin A/C. It will also be important to validate the role of Lap2α in regulating the other MRTF family members, MRTF-B and myocardin.

While the loss of Lap2α led to decreased binding of MRTF-A to the SRF target genes, our ChIP-seq data surprisingly revealed a higher number of MRTF-A peaks in the Lap2α KO cells compared to WT cells in starved conditions (**Fig S4A**). This result might be explained by increased nuclear localization of MRTF-A in starved KO cells that we showed in our localization experiments (**Fig 3**), and might reflect binding of MRTF-A to lower affinity binding sites on chromatin. Motif analysis revealed that MRTF-A peaks were enriched for motifs of other transcription factors, such as ZBTB33 (**Table S4**). Further experiments are required to elucidate the role of MRTF-A in SRF-independent transcription, and its cooperation with other transcription factors. Here it is also interesting to note that our co-immunoprecipitation experiments demonstrated the requirement for the MRTF-A SAP-domain in binding to Lap2α (**Fig 6I**). The role of the SAP domain in MRTF-A function is still somewhat elusive. Its structure points to a possible interaction with DNA similarly to other SAP-domain containing proteins (Aravind and Koonin 2000) and at least a set of specific genes associated with breast cancer have been suggested to be regulated by the MRTF SAP-domain, in SRF-independent manner (Gurbuz et al. 2014). Whether the interaction with Lap2α plays a role here awaits further studies.

Taken together, our research describes a novel mechanism to regulate the expression of MRTF-A/SRF target genes by Lap2α. In the future, it will be interesting to study further the biological relevance of this regulatory interaction in the context of both normal development and disease, and to explore further the molecular mechanisms involved.

## Material and methods

### Antibodies

| name | company/source | application |
| --- | --- | --- |
| RNA polymerase II CTD | Abcam ab 5408 | ChIP-seq |
| MRTF-A | home-made rabbit antiserum provided by G.Posern (Descot et al, 2009) | ChIP-seq |
| Lap2α | mouse monoclonal antibody Gesson et al., 2016) | ChIP-seq |
| SRF | Cell Signaling Rabbit mAB #5147 | ChIP-seq |
| IGG | Santa Cruz sc-2027 | ChIP-seq |
| MRTF-A | Santa Cruz G-8 sc-390324 1:50 | IF |
| Alexa-Fluor-488-conjugated anti-mouse | Invitrogen A21155 1:500 | IF |
| Alexa-Fluor-488-conjugated anti-rabbit | Invitrogen A11008 1:500 | IF |
| HRP-conjugated Flag M2 | Merck A8592 1:10000 | WB |
| HRP-conjugated HA | Merck H6533 1:7500 | WB |
| Lap2α | Immuquest IQ175 1:2000 | WB |
| Lap2 | Abcam ab 185718 1:1000 | WB |
| HRP-conjugated His | Merck A7058 1:5000 | pull-down |

### Plasmids

Details of the plasmid cloned for the study are available on request.

Plasmids used in the study: MRTF-A-GFP-N3, MRTF-A-2HA-N3, MRTF-A-2Flag-N3, MRTF-A-C471(Miralles et al. 2003), MRTF-A-ΔN-2HA-N3, MRTF-A-ΔQ-2HA-N3, MRTF-A-ΔSAP-2HA-N3, MRTF-A SAPmut, MRTF-A-ΔLZ-2HA-N3, RPEL-2HA-N3, MRTF-A-pDest10, Lap2α-2HA-C1, Lap2α-2Flag-C1, pET41a-3CΔ-Lap2α-CTD, reporter plasmids were p3DA.luc (Geneste et al. 2002), pRL-TK (Promega). For the design of MRTF-A SAPmut, we first analyzed the structure of MRTF-A SAP domain using Phyre^2^ web portal (Kelley et al. 2015). Using structure of MRTF-A SAP domain (https://www.rcsb.org/structure/2KVU) as a reference, Phyre^2^ identified several SAP domain-containing proteins, and we chose three of them characterized by high confidence (>98%) and high sequence identity (47-100%) of their SAP domains: ERI1 (human), SIZ1 (rice), THO1 (yeast). We did alignment of amino acid sequences and compared it with the data from Aravind and Koonin (Aravind and Koonin 2000), who have described SAP domain as a putative DNA-binding motif. Although the most conserved residues were represented by five leucines, we decided not to mutate them because they are required to maintain correct folding of SAP domain helixes. We chose four amino acid residues, which appeared the most conserved among charged residues in SAP domains of analyzed proteins and replaced them with non-charged serine (K451S, R457S, K465S and R471S) in MRTF-A.

### Cell lines and transfections

NIH 3T3 cell line were obtained from the Treisman lab. Tetracycline-inducible MRTF-A-GFP-expressing NIH 3T3 cell line (R332) is described in (Vartiainen et al. 2007). Mouse dermal fibroblast (MDF) cell line with knock-out of Lap2α and the corresponding wild type MDF cell line were derived from Lap2α knockout mice (Naetar et al. 2008). All cell lines were cultured in Dulbecco’s modified Eagle’s medium (DMEM, Lonza) supplemented with 10% Fetal Bovine Serum (FBS; GIBCO), 100 units/ml Penicillin and 0.1mg/ml Streptomycin (Thermo Fisher Scientific) and maintained in humidified 95% air/5% CO2 incubator at +37°C. In addition, the R332 cell line was supplemented with 5 μg/ml Blasticidin (Invivogen) and 250 μg/ml Zeocin (Invivogen). Tetracycline (1μg/ml) was added to the growth medium to induce the expression of MRTF-A-GFP.

For siRNA transfections, NIH 3T3 cells were plated onto 6-well tissue culture plate at a density of 25000 cells per well one day prior to transfection. On the next day, cells were transfected with 20 nmol siRNA (mouse Lap2: 5’-GAUGUGACAGAGCUCUCUA predesigned from Sigma; mouse Lap2α: 5’-AGUCCAGCUAUCAAGAUUC; mouse Lap2β: 5’ – GAGUACUCCCAUAGCUGAA; negative control: AllStars negative control from Qiagen) using Interferin transfection reagent (Polyplus transfection) according to manufacturer’s protocol. On day 4, cells were re-transfected with siRNAs and DNA constructs if needed (for the Luciferase assay, see below), and the following day processed for western blotting, RNA extraction or luciferase assay.

For Western blotting analysis, cells were washed with 1x PBS and lysed in 1x Laemmli buffer. For RNA extraction and RT-qPCR analysis, re-transfected cells maintained in low-serum (0.3% FBS) DMEM for 24 hours. After starvation, cells were stimulated with serum (15%) for 45 min and harvested for RNA extraction.

### Luciferase assay

MDF or NIH cells were plated onto a 24-well plate. Next day MDF cells were transfected with SRF reporter (8 ng) and reference reporter pTK-RL (20 ng), with or without normalization control SRF-VP16 (20 ng) by using JetPrime transfection reagent. NIH cells were first transfected with siRNA as above, on day 4, re-transfected with p3DA.luc and pTK-RL plasmids. Cells were maintained in DMEM containing 0.3% FBS for 24 h and stimulated with serum (15%). After 7 h of stimulation, cells were harvested and analysed with the Dual-Luciferase reporter assay system (Promega) and a Multimode Plate Reader EnSpire (PerkinElmer), according to manufacturer’s instructions. For data analysis, the activity of firefly luciferase was normalized to the renilla luciferase activity and SRF-VP16 was set to 100.

### Immunostaining and microscopy

For microscopy, cells on coverslips were fixed with 4% paraformaldehyde for 15 min, washed four times with PBS and permeabilized for 5 min with 0,5% Triton X-100 (Merck) in PBS. For antibody staining, permeabilized cells were blocked with a blocking buffer (1% BSA, 1% Gelatin, 10% FBS) for 30 min and incubated with primary antibody for 1 h at RT. Coverslips were washed three times with PBS and incubated with Alexa Fluor-conjugated secondary antibody for 1 h with or without DAPI (4′,6-diamidino-2-phenylindole). Coverslips were washed three times with PBS and mounted in Moviol 4-88 (Merck). Wide-field fluorescence microscope Leica (Leica, Welzlar, Germany) DM6000 with HCXPL APO 63x/1.40-0.60 oil objective and confocal Zeiss LSM700, 63x/1.3 objective and ZEN software were used to image the samples. The image files were processed with MIB software(Belevich et al. 2016).

### RNA seq

For RNA seq, NIH 3T3 cells were plated onto 10 cm plates. The following day, cells were transfected with siRNA as above (Lap2: 5’-GAUGUGACAGAGCUCUCUA from Sigma; negative control: AllStars negative control from Qiagen). Total RNA was extracted with a Nucleospin RNA kit from Macherey-Nagel according to the manufacturer’s protocol from triplicates of control and Lap2-depleted samples. Libraries were prepared for Illumina NextSeq 500 using Ribo-Zero rRNA Removal Kit (Illumina) and the NEBNext Ultra Directional RNA Library Prep at the Biomedicum Functional Genomics Unit (FuGU) according to the manufacturer’s protocols. RNA-seq data sets were aligned using TopHat2(Kim et al. 2013) (using Chipster software (Kallio et al. 2011)) to version mm10 of the mouse genome with the default settings. Counting aligned reads per genes were performed with HTSeq (Anders et al. 2015). Differential expression analysis was performed with DESeq (Love et al. 2014). List of the transcribed genes was based on the aligned reads count cutoff >1 from the RNA-Seq data from control vs Lap2-depleted NIH 3T3 cells. Gene ontology was performed using DAVID (Huang da et al. 2009a; Huang da et al. 2009b).

### ChIP-seq

For chromatin immunoprecipitation (ChIP), NIH 3T3 or MDF cells were grown on 15 cm plates. Confluent cells (5×10⁶ cells) were fixed in 1% paraformaldehyde/PBS for 10 min at RT, crosslinking was stopped by adding cold glycine to a final concentration of 0.125 M for 5 min, followed by harvesting in 500 ml of ice-cold PBS and spinning at 1000g for 5 min at +4°C. Cell pellets were lysed in 300 μl of RIPA buffer and sonicated with Bioruptor (Diagenode; number of cycles = 15, power = HIGH, ON = 30 sec, OFF = 30 sec). IPs were carried out with 5 μg antibody overnight at 4°C in a rotating wheel. The immuno-complexes were collected with 50 μl of protein A sepharose (17-0780-01, GE Healthcare) at 4 °C for two hours with rotation. The beads were pelleted by centrifugation at 4 °C for 1 min at 500g and washed sequentially for 5 min on rotation with 1 ml of the following buffers: low-salt wash buffer (RIPA) (10 mM Tris–HCl (pH 8.0), 0.1% SDS, 1% Triton X-100, 1 mM EDTA, 140 mM NaCl, 0,1% sodium deoxycholate), high-salt wash buffer (10 mM Tris–HCl (pH 8.1), 0.1% SDS, 1% Triton X100, 1 mM EDTA, 500 mM NaCl, 0,1% sodium deoxycholate) and LiCl wash buffer (10 mM Tris– HCl (pH 8.1), 0.25 mM LiCl, 0,5% IGEPAL CA-630, 0,5% sodium deoxycholate, 1 mM EDTA). Finally, the beads were washed twice with 1 ml of TE buffer (10 mM Tris–HCl (pH 8.0), 1 mM EDTA). Chromatin was eluted in 150 μl of 1% SDS in TE buffer. The cross-linking was reversed by adding NaCl to final concentration of 200 mM and incubating at 65 °C overnight. The eluate was treated with Proteinase K and the DNA was recovered by extraction with phenol/chloroform/isoamyl alcohol (25/24/1, by vol.) and precipitated with 0.1 volume of 3 M sodium acetate (pH 5.2) and two volumes of ethanol using glycogen as a carrier. ChIP libraries were prepared for Illumina NextSeq 500 using NEBNext ChIP-Seq DNA Sample Prep Master Mix Set for Illumina (NEB E6240) and NEBNext^®^ Multiplex Oligos for Illumina^®^ (Index Primers Set 1) (NEB E7335) according to the manufacturer’s protocols. Sequencing was performed with NextSeq500 at Biomedicum Functional Genomics Unit (FuGU). ChIP-Seq data sets were aligned using Bowtie2 (using Chipster software (Kallio et al. 2011)) to version mm10 of the mouse genome with the default settings. Peak calling was performed with MASC2 (Zhang, Y. et al. 2008). To visualize and present ChIPseq data, we used Integrative Genomics Viewer, IGV (Robinson et al. 2011) and EaSeq (http://easeq.net)(Lerdrup et al. 2016). We used MEME-ChIP to perform comprehensive motif analysis (including motif discovery) (Bailey et al. 2009).

### Data access

ChIP-seq and RNA-seq data are available under Gene Expression Omnibus accession number GSE159371.

### Single cell tracking assay

MDF cells were plated in 12-well plate (3000-4000 cells/well) in 10% FBS DMEM. On the next day, medium was replaced to 0.3% FBS DMEM, and cells were maintained under starved conditions for 24 hours. After starvation, cells were stimulated with serum (15%) and immediately transferred into Cell-IQ Fluorescence imaging system (Chip-Man Technology) and monitored for 16 hours at +37°C and CO2 flow 25-30 mL/min. The images were taken with 20 min intervals. Images were stitched using wound healing assay tool of MIB software(Belevich et al. 2016); the stitched images were downsampled two times and saved as 3D-TIF stacks. The cell tracking was done using TrackMate plugin of Fiji (Tinevez et al. 2017). To make unbiased selection of cells, a grid (100 × 100 μm) was applied to datasets and the cells closest to the grid line intersections were selected for tracking. The average speed of each cell was calculated and the results for all cells were plotted using violin plot function (Hoffmann 2015) in Matlab (Mathworks Inc, MA).

### Co-IP assays

NIH 3T3 cells (1 × 10^6^) were plated on 10 cm dishes and transfected with the appropriate HA/FLAG-tagged constructs (3 μg each plasmid) using Jet Prime transfection reagent. Twenty –four hours later, cells were harvested in IP buffer (0.5% Triton X-100, 50 mM Tris-HCl pH 7.5, 150 mM NaCl, protease inhibitor cocktail (Sigma-Aldrich)) and the lysates were cleared by centrifugation. Cleared lysates were then subjected to EZview Red Anti-HA Affinity Gel or Anti-FLAG M2 Affinity Gel (Sigma-Aldrich), incubated 3 h on rotation at +4 °C and washed three times with IP buffer without Triton X-100. Bound proteins were eluted with 1 × SDS–PAGE loading buffer, boiled for 5 min, separated in 4-20% SDS–PAGE (BioRad) and electroblotted to nitrocellulose membrane. The membrane was probed with indicated antibodies.

### Real-time quantitative PCR

NIH or MDF cells were plated on 35 mm dishes. On the next day, the media was changed to fresh growth media containing 0.3% FBS. Cells were maintained in low-serum conditions for 24 hours. On day 3, cells were stimulated with 15% FBS for 45 min and total RNA was extracted using Nucleospin RNA II kit according to the manufacturer’s protocol (Macherey-Nagel). Five hundred micrograms of total RNA wasused for complementary DNA synthesis using Thermo Scientific RT–PCR kit (Thermo Scientific). Quantitative PCR was carried out using the Bio-Rad CFX machine (Bio-Rad) and SYBR green qPCR reagent (Thermo Scientific). Gene specific primers are listed below:

Gapdh_fw: 5′-TGCACCACCAACTGCTTAGC-3′

Gapdh_rev: 5′-GGCATGGACTGTGGTCATGAG-3′

Acta2 fw: 5′-ACTGGGACGACATGGAAAAG-3′

Acta2 rev: 5′-GTTCAGTGGTGCCTCTGTCA-3′

Vcl fw: 5′-CTTTGTGCAGGCAAGGAACG-3′

Vcl rev: 5′-GCTGCATTCTCCACTTTGGC-3′

Tagln fw: 5′-CTATGAAGGTAAGGATATGGC

Tagln rev: 5′-TCTGTGAAGTCCCTCTTATG

Egr1 fw: 5′-CAGAGTCCTTTTCTGACATC

Egr1 rev: 5′ – GAGAAGCGGCCAGTATAG

Egr3 fw: 5′-CCACCTCACCACTCACATCC

Egr3 rev: 5′ – CTTGAGGTGGATCTTGGCGT

Fos fw: 5′-TACTACCATTCCCCAGCCGA

Fos rev: 5′ - GCTGTCACCGTGGGGATAAA

Relative expression levels were calculated by the comparative *C*T method, normalizing to the Gapdh cDNA: 2^−C^_T_ (target)/2^−C^_T_ (Gapdh).

### Expression and purification of MRTF-A protein

Recombinant mouse His-MRTF-A was expressed by using MultiBac baculovirus expression system. MRTF-A-pDest10 plasmid was transformed into DH10MultiBac cells, recombinant bacmids were prepared and the presence of the correct construct verified by PCR. The verified bacmid was transfected into Sf21 cells using Fugene HD transfection reagent (Promega). The obtained viral stock was amplified and used for protein expression. The expression of His-MRTF-A was done by infecting cells at the density of 2,0 × 10^6^ cells/ml with the baculoviral stock at a multiplicity of infection (MOI) of 1. The cells were collected after 96 hours and the pellets were snap frozen and stored at −80°C. The expression was verified by Western blot. For purification, the cell pellets were resuspended in lysis buffer (25 mM Tris-HCl pH 8.0, 10 mM NaCl, 1 mM MgCl_2_, 2 mM B-mercaptoethanol, 0,2 % Igepal) with protease inhibitors (Roche) and benzonase nuclease (Sigma-Aldrich) and lysed with EmulsiFlex-C3 (AVESTIN) with gauge pressure ~15 000 psi for 10 min. Immediately after lysis, the NaCl concentration was adjusted to 200 mM and 10 mM imidazole was added. Cell extract was incubated for 10 min at +4°C and clarified by centrifugation at 39 000 g for 30 min and immediately processed to metal (Ni^2+^) affinity purification using Ni^2+^-NTA agarose beads (Qiagen) equilibrated with lysis buffer. Beads were incubated with extract for 2 h on rotation at +4°C and washed three times with buffer A (25 mM Tris-HCl pH 8.0, 300 mM NaCl, 2 mM B-mercaptoethanol) and two times with buffer A with addition of 20 mM imidazole. The last wash was conducted in a disposable 10 ml Poly-prep column (Bio-RAD) and the protein was eluted from the beads with elution buffer (25 mM Tris-HCl pH 8.0, 100 mM NaCl, 500 mM imidazole). The protein was further purified with gel filtration column Superdex 200 HiLoad 16/60 (Pharmacia), which was equilibrated with buffer B (50 mM Tris-HCl pH 8.0, 100 mM NaCl, 1 mM DTT, 0,5 mM EDTA, 10 % glycerol). Fractions were analyzed by SDS-PAGE and those containing His-MRTF-A were concentrated and stored at −80°C.

### Expression of Lap2α-CTD

The coding region corresponding to murine Lap2α residues 459-693 was cloned to pET 4.1-3CD. *E.coli* cells expressing His-GST-Lap2α-CTD were grown at 37°C for 2-3 hours until OD_6 0 0_ reached 0.6-1.0 and then induced with IPTG to a final concentration of 1mM. Then cells were cultivated for 3 hours at 37°C, harvested by centrifugation at 4000 rpm 3 min, frozen and stored at −80°C.

### Pull-down assay

His-GST (10 μl) and His-GST-Lap2α-CTD (100 μl) *E.coli* lysates were immobilized on 50 μl of Protino 4B glutathione agarose (Macherey-Nagel) equilibrated with binding buffer 1 (25 mM Tris-HCl pH 7.5, 150 mM NaCl, 1 mM DTT) in a total reaction volume of 300 μl in binding buffer 1 for 2 hours on rotation at +4°C. Beads were washed once with binding buffer 1, three times with binding buffer 2 (25 mM Tris-HCl pH 7.5, 500 mM NaCl, 1 mM DTT) and once with binding buffer 1. After the washes, beads were incubated with 1,5 μM purified recombinant His-MRTF-A in binding buffer 1 for overnight on rotation at +4°C. After incubation the beads were washed three times with binding buffer 1 and bound proteins were eluted with 1xSDS-PAGE loading buffer. Samples were boiled for 5 min and the proteins were separated in 4-20% Mini-PROTEAN TGX Precast Protein Gel (Bio-RAD) and electroblotted to nitrocellulose membrane. The membrane was probed with anti-His antibody to assess the association of His-MRTF-A with GST-fusions.

### Statistical analysis

Statistical analyses were performed in Excel or XLSTAT by two-tailed Student’s *t*-test, with two-sample unequal variance for the data conformed to normal distribution (SRF reporter activity, expression of SRF target genes, MRTF-A localization). Non-parametric Kolmogorov-Smirnov test, with the significance level of 0.05, was applied to Single cell tracking assay data, because the data did not conform to normal distribution.

## Acknowledgements

We thank Paula Maanselkä for excellent technical assistance. Imaging was performed at Light Microscopy Unit (LMU), which is supported by Helsinki Institute for Life Science (HiLIFE) and Biocenter Finland. This work was supported by grants (to MKV) from Sigrid Juselius foundation, Academy of Finland, Jane and Aatos Erkko foundation, Finnish Cancer foundation and HiLIFE.

## Author contributions

ES, MS and MKV conceived and planned out the experiments. ES, AP and SK performed the experiments. ES and MS analyzed the results. GP and RF provided reagents and intellectual input into the project. ES, MS and MKV wrote the manuscript based on input from GP and RF. MKV supervised and obtained funding for the project.

## Supplementary figures

**Figure S1.**
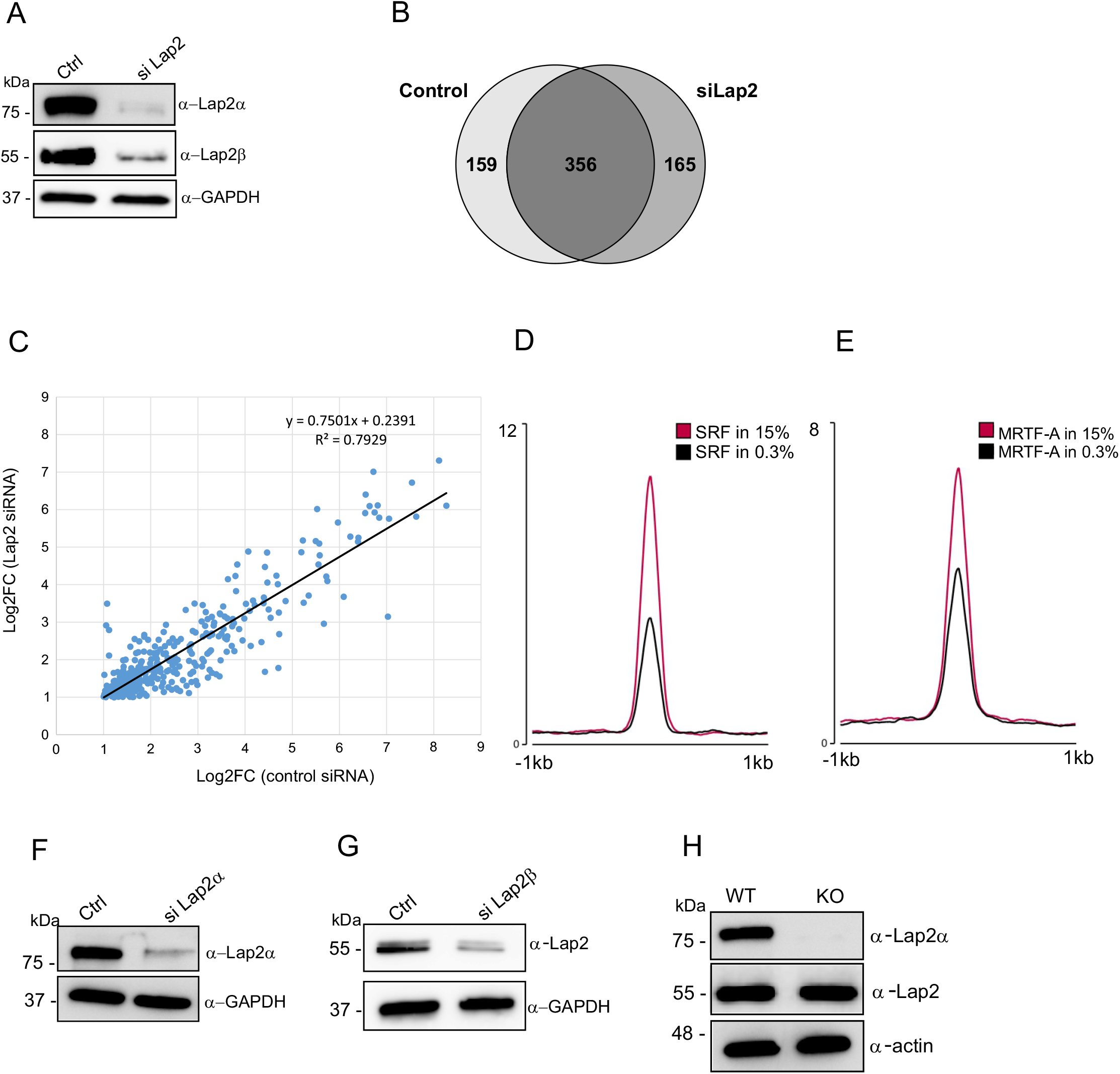
Related to figure 1. **(A)** Western blot showing depletion of Lap2α and Lap2ß in NIH 3T3 with siRNA against total Lap2. Used siRNAon the top, molecular weight markers on the left, and antibody used detection on the right. **(B)** Venn diagram showing overlap of serum-responsive genes in control and Lap2-depleted NIH 3T3 fibroblasts. **(C)** Differential expression scatter plot showing that serum-activated genes have higher activation in control cells than in Lap2-depleted NIH 3T3 fibroblasts. **(D)** SRF and **(E)** MRTF-A enrichment on the SRF peaks identified in serum-stimulated conditions in NIH 3T3 fibroblasts. **(F)** Western blot showing depletion of Lap2α in NIH 3T3; data shown as in A. **(G)** Western blot showing depletion of Lap2b in NIH 3T3; data shown as in A **(H)** Western blot showing Lap2α and Lap2b protein amounts in Lap2α WT and KO mouse dermal fibroblasts.

**Figure S3.**
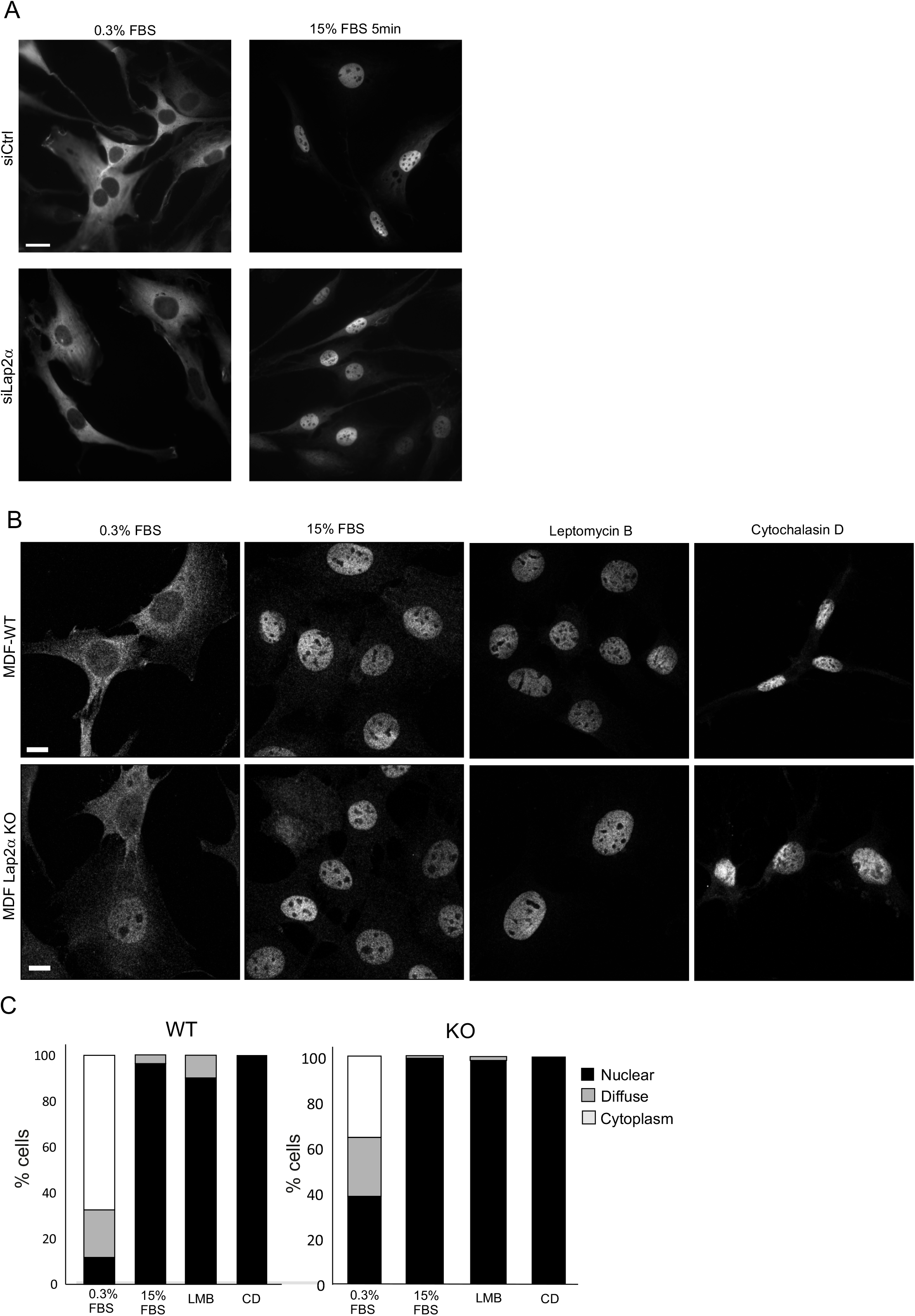
Lap2α is not required for MRTF-A nuclear localization; related to figure 3. **(A)** Localization of MRTF-A-GFP in serum-starved (0.3% FBS), and serum-stimulated (15% FBS) NIH 3T3 fibroblasts transfected with control (upper panel) or Lap2α (lower panel) siRNAs. Scale bars 10 μm. **(B)** Localization of endogenous MRTF-A in serum-starved (0.3% FBS), serum-stimulated (30 min, 15% FBS), Leptomycin B-treated (30 min, 20 nm) and cytochalasin D (CD)-treated (30 min, 2μm) Lap2α WT (upper panel) and KO (lower panel) MDF cells. Scale bars 10 mm. **©** Quantification of endogenous MRTF-Alocalization in Lap2α WT (left) and KO (right) MDFs. Number of cells: N=154 WT 0.3%; N=141 WT 15%; N=158 WT LMB; N=129 WT CD; N=195 KO 0.3%; N=178 KO 15%; N=231 KO LMB; N=52 KO CD

**Figure S4.**
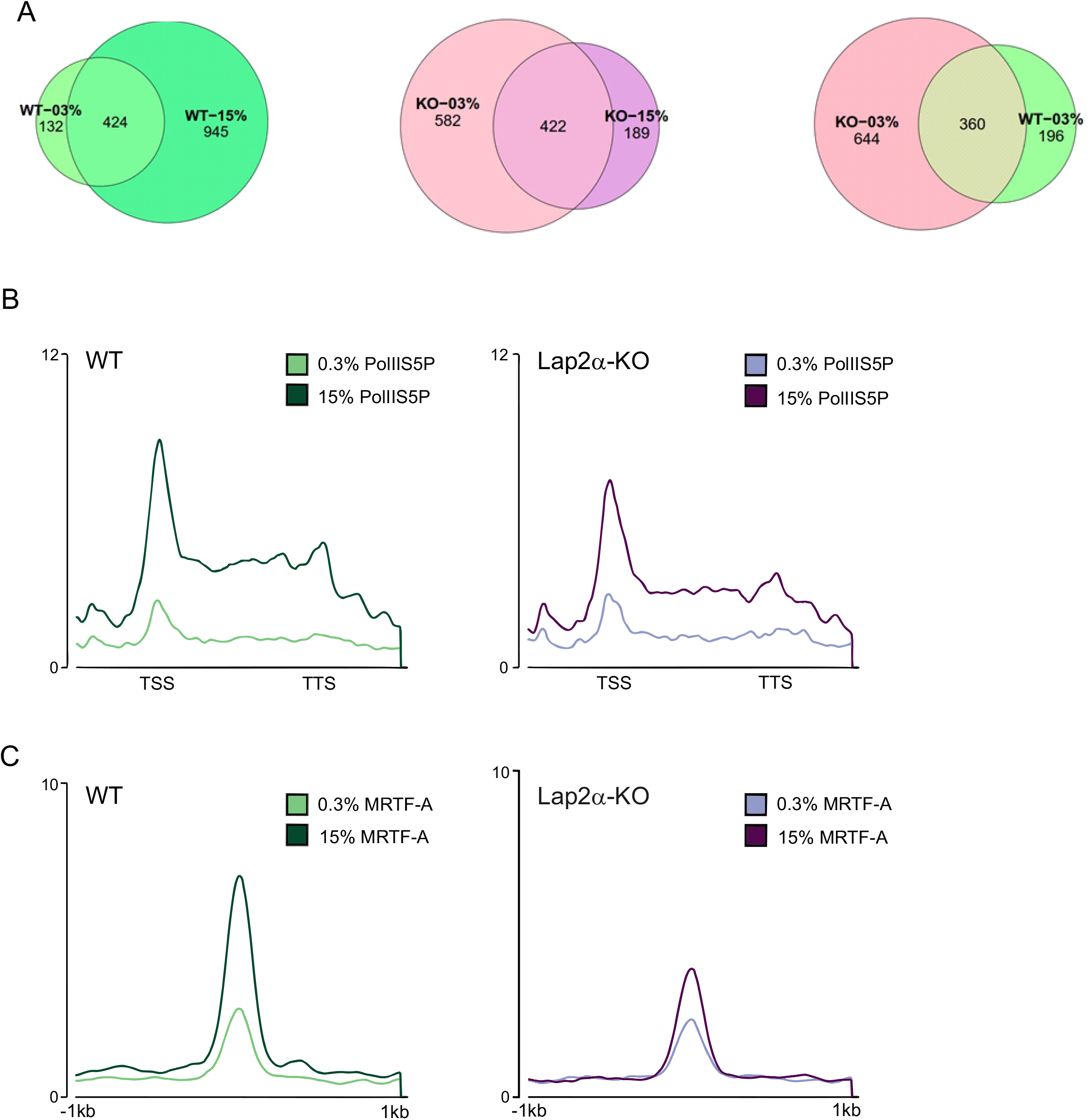
Chromatin-binding of MRTF-A and Pol II in Lap2α WT and KO cells, related to figure 4. **(A)** Venn diagrams showing overlaps of MRTF-A peaks enriched with SRF motif in Lap2α WT and KO MDF in serum-starved (0.3% FBS) and serum-stimulated (15% FBS) conditions. **(B)** Pol II S5P enrichment on selected 36 genes with MRTF-A peaks enriched with SRF motif (Table S3.2) in Lap2α WT and KO MDF **C)** MRTF-A enrichment on selected 36 genes with MRTF-A peaks enriched with SRF motif (Table S3.2) in Lap2α WT and KO MDF

**Figure S5.**
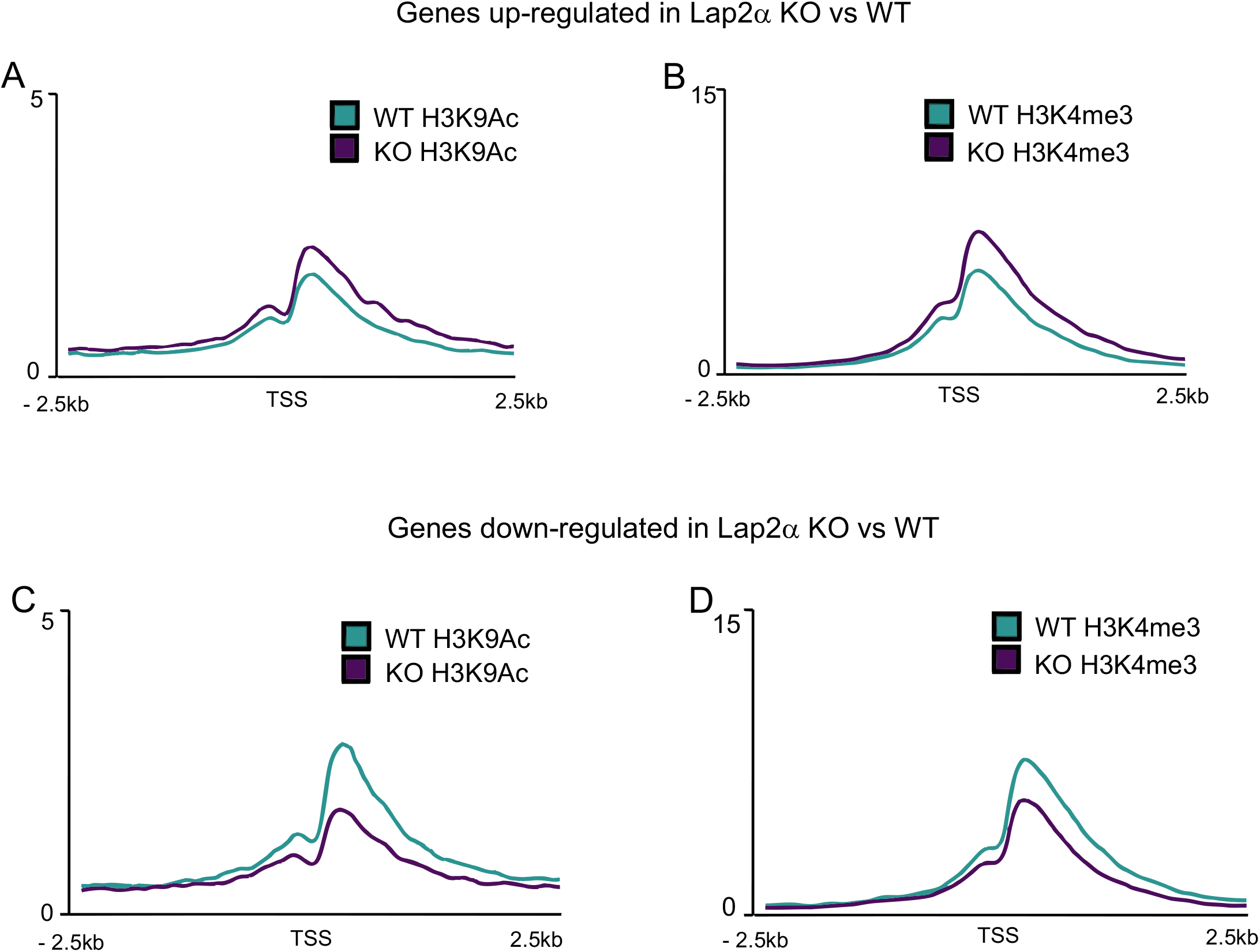
Distribution of active histone marks (H3K4me3 and H3K9Ac) on up- and down-regulated genes in Lap2α KO vs WT MDF. Reanalyzed from Gesson et al, 2016. **(A,B)** Enrichment of active histone marks H3K9Ac (A) and H3K4me3 (B) on TSS of genes up-regulated in Lap2α KO fibroblasts is higher in KO cells. **(C,D)** Enrichment of active histone marks H3K9Ac (C) and H3K4me3 (D) on TSS of genes down-regulated in Lap2α KO fibroblasts is higher in WT cells.

**Figure S6.**
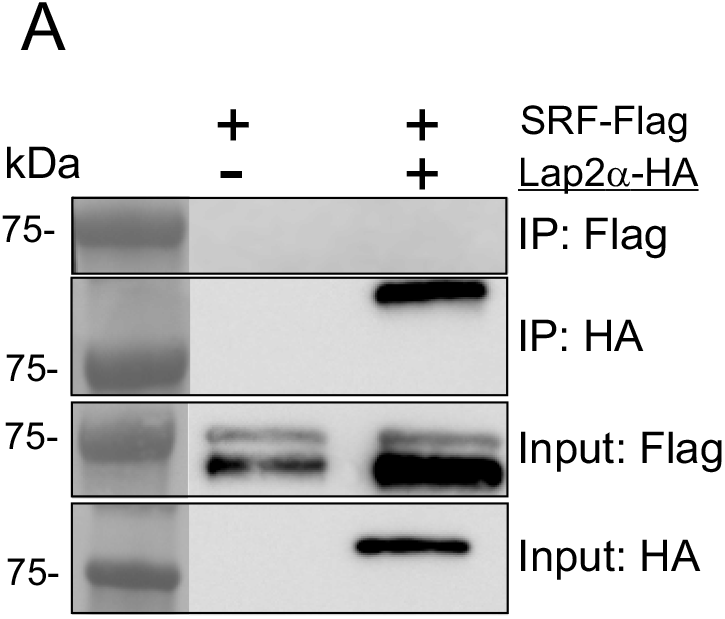
Lap2α does not interact with SRF. (**A**) HA-tagged Lap2a does not co-immunoprecipitate Flag-tagged SRF. Transfected constructs are shown above the blots with the construct used as a bait underlined, protein weight markers on the left and the sample (immunoprecipitate or input) with the antibody used in Western on the right.

## Supplementary Tables

**Table S1. RNA-seq data**
  **S1.1** RNA-seq in Lap2-depleted NIH 3T3 cells
  **S1.2** Upregulated genes (515) in control samples after serum stimulation.
  **S1.3** Gene ontology enrichment analysis of genes downregulated in Lap2-depleted NIH 3T3 cells
**Table S2.Total amount of MRTF-A peaks in Lap2α WT and KO MDF cells**
**Table S3. MRTF-A peaks with SRF motif**
  **S3.1** MRTF-A peaks with SRF motif in Lap2α WT MDFs in 15% FBS
  **S3.2** 36 selected genes contained MRTF-A peaks with an SRF binding motif on their promoters
**Table S4. Data of MEME-ChIP analysis**

